# Juggling offsets unlocks RNA-seq tools for fast and Scalable differential usage, Aberrant Splicing and Expression Retrieval

**DOI:** 10.1101/2023.06.29.547014

**Authors:** Alexandre Segers, Jeroen Gilis, Mattias Van Heetvelde, Davide Risso, Elfride De Baere, Lieven Clement

**Affiliations:** *Department of Applied Mathematics, Computer Science and Statistics, Ghent University, Ghent, Belgium; Center for Medical Genetics Ghent, Ghent University and Ghent University Hospital, Ghent, Belgium; Data Mining and Modeling for Biomedicine, VIB Flemish Institute for Biotechnology, Ghent, Belgium; Bioinformatics Institute Ghent, Ghent University, Ghent, Belgium.; Department of Biomolecular Medicine, Ghent University, Ghent, Belgium; Department of Statistical Sciences, Universiy of Padova, Padova, Italy

**Keywords:** RNA-sequencing, Aberrant expression, Aberrant splicing, Differential transcript usage, Offsets, Normalisation

## Abstract

RNA-seq data analysis relies on many different tools, each tailored to specific applications and coming with unique assumptions and limitations. Indeed, tools for differential transcript usage or rare disease diagnosis through splicing and expression outliers, either lack performance, discard information, or do not scale to large datasets. We show that replacing normalization offsets unlocks bulk RNA-seq tools for differential usage and aberrant splicing, providing a single framework for various short- and long-read applications. We then introduce saseR, a tool for prioritizing expression and usage outliers that is much faster than state-of-the-art methods, and significantly outperforms these for aberrant splicing detection.

## 1 Background

RNA-seq technologies revolutionized biological and biomedical research by providing a comprehensive, quantitative, and unbiased view on the transcriptional landscape underpinning complex biological processes and disease. As the technologies became mature, the key challenges switched from data generation to data analysis that has to scale to the ever-increasing data volumes in contemporary RNA-seq based studies. In Figure 1 we give an overview of different research hypotheses that are commonly assessed with bespoke RNA-seq data analysis workflows and tools to discover disease-related mechanisms.

**Fig. 1.**
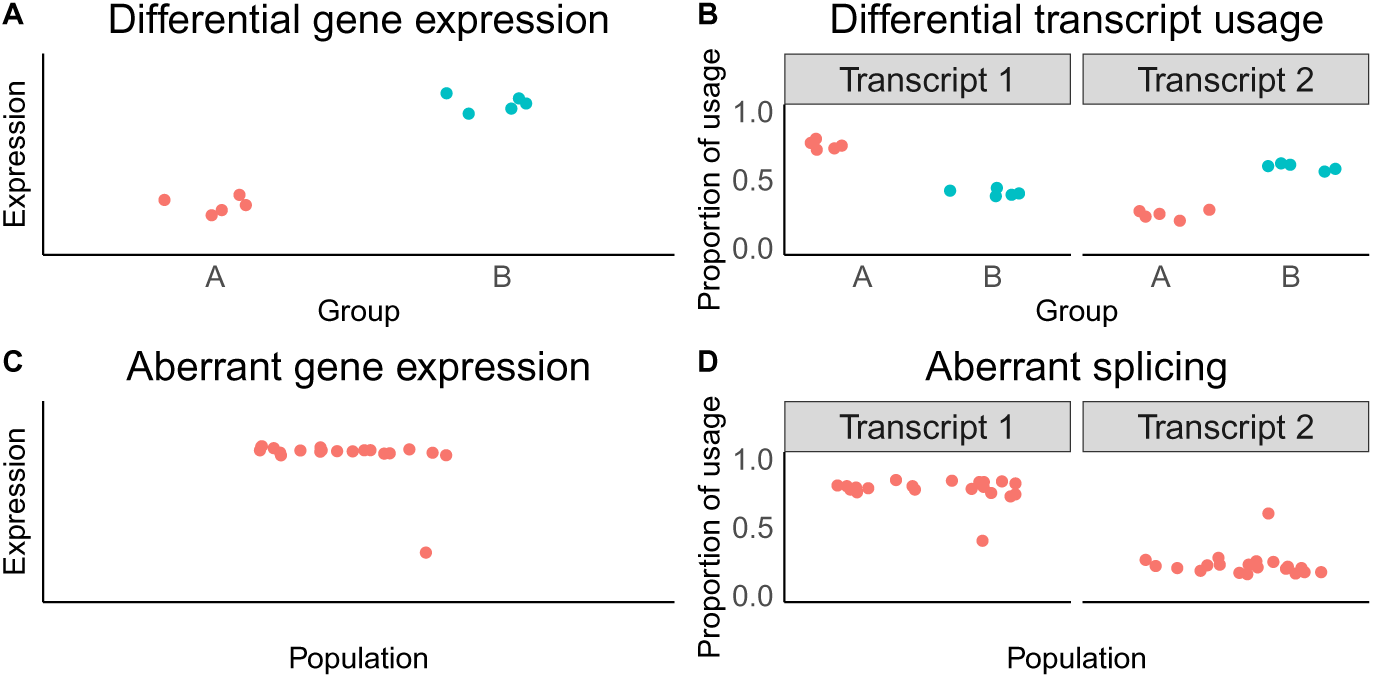
Different analysis methods used to discover disease-related mechanisms. Differential gene expression methods compare the average expression between two groups (panel A). Differential transcript usage methods compare the proportion of the transcript expression relative to its total gene expression, between two groups (panel B). Outlier or aberrant analyses search for an extreme value in one patient, rather than comparing two groups, due to the lack of replicates. This can be done for both expression analysis (panel C as shown in [1]) and splicing or usage analysis (panel D).

A first task to unravel biological processes and disease is the prioritization of differentially expressed genes or transcripts. Indeed, bulk differential gene expression (DGE) tools such as edgeR and DESeq2 are well established, are flexible to model data with complex designs and scale to the data volumes of contemporary studies involving hundreds of patients.

The dysregulation of splicing has been shown to be another important driver of disease [2–5], including frontotemporal dementia, Parkinsonism and spinal muscular atrophy, and is a well-known hallmark of cancer [6]. Prioritizing genes and transcripts that are associated with the dysregulation of splicing has become possible by widespread adoption of expression quantification through pseudo-alignment [7, 8], which enables fast and accurate quantification of expression at the transcript level. Again advances in the data analysis have become crucial in this context. Indeed, novel tools that can assess differential transcript usage (DTU), i.e. change in relative usage of transcripts/isoforms within the same gene, have become key to further unravel the molecular basis of disease. However, bespoke tools have been reported to be suboptimal because they either do not scale to the ever increasing data volumes or cannot account for multiple covariates, which are required to model the data of multifactorial designs and to correct for batch effects [9].

More recently, RNA-seq technologies have also been used to unravel the molecular basis of Mendelian diseases, which are estimated to impact 350 million people world-wide [10, 11]. For these individuals, a diagnostic rate of 15-75% is currently achieved by using whole genome sequencing or whole exome sequencing to identify the underlying pathogenic variants [12–15]. Although most of these variants are within coding regions, there is increasing evidence that the diagnostic rate can be further improved by discovering variants in intronic regions and in other non-coding regions that contribute to different splicing and/or impaired transcriptional regulation [16]. However, using default DGE and DTU bulk RNA-seq workflows in the context of rare diseases is not possible because different patients suffer from mutations in different genes. Therefore, replicates are lacking, which renders testing each sample against the rest of the cohort statistically invalid. In this regard, new data analysis tools have been developed to pick up genes with aberrant expression [1, 17] or splicing [18–21] in each sample of a diseased cohort. Hence, the data analysis is directed towards expression or splicing outliers in each subject using count-based outlier tests.

Although analysis tools for DGE such as edgeR [22] and DESeq2 [23] are well established and fast, fewer tools have been developed for DTU, aberrant expression and aberrant splicing, and these are not yet as scalable. For DTU, DEXSeq [24] and diffspliceDGE of edgeR are often used, but the former becomes slow for large data compendia while the latter has lower power to detect DTU events [9]. In this regard, we developed satuRn [9], a scalable alternative for DTU analyses. However, satuRn builds on a quasi-likelihood modeling framework in which only the first two moments of the distribution are modelled without specifying the full distribution. Therefore, it cannot be used for detecting expression or aberrant splicing outliers. For the detection of aberrant expression, OUTRIDER [1] uses a negative binomial autoencoder to control for latent confounders and uses count-based outlier tests to prioritise genes. However, computing the negative binomial autoencoder is slow. To overcome this computational burden, Outsingle [17] uses a log-normal approximation, but therefore ignores the discrete count nature of bulk RNA-seq data, and can moreover not control for known confounders. For aberrant splicing analyses, LeafcutterMD [18] and SPOT [19] consider outlier detection using a Dirichlet-Multinomial distribution. However, these tools cannot control for confounders (e.g. batch effects) in clinical datasets. To improve upon this, the state-of-the-art method for aberrant splicing detection in bulk RNA-seq, FRASER [20, 21], uses a similar approach as OUTRIDER by using a beta-binomial autoencoder to control for latent confounders, which again comes with a considerable computational burden. Moreover, FRASER rejects information by only using junction-reads, ignoring exonic and intronic bin reads.

Hence, in the current bioinformatics landscape, the data analysis to unravel the molecular basis for disease based on RNA-seq data involves a plethora of different tools, that each have their own assumptions, that are often restricted to experiments with simple designs, and that often do not scale to the ever increasing data volumes. In this contribution, we provide data analysts with a unified workflow for differential expression analysis, differential usage analysis, and detecting expression and usage outliers, that is scalable and flexible to model the ever increasing data volumes of contemporary RNA-seq studies with complex designs and multiple batch effects. Particularly, we start from well established bulk RNA-seq DGE workflows and show how juggling offsets can effectively unlock these for fast and scalable differential usage and aberrant splicing analyses. Indeed, by replacing the conventional offset for library size in DESeq2 or edgeR transcript or exon level analyses with the logarithm of the total gene count, the parameters of the mean model enable us to directly estimate the average transcript or exon usage, respectively. Further, by estimating the means and gene-specific dispersion of the negative binomial (NB) distribution, one can evaluate how likely it is that the observed read-counts are generated from the estimated NB distribution, extending conventional DGE frameworks to detect aberrant expression and splicing outliers in the context of rare diseases. We also evaluate workflows on different ASpli [25] counts, which are specifically developed for the quantification of differential splicing, i.e. bin and junction counts, combined with the appropriate offsets for inferring aberrant splicing. Finally, we develop a fast algorithm for parameter estimation to assess aberrant expression and splicing that scales better to the large number of latent covariates that are typically needed in studies on rare disease with large cohorts, which we implemented and distributed in a Bioconductor R package called saseR (Scalable Aberrant Splicing and Expression Retrieval). In simulation and real case studies we show how saseR, vastly outperforms existing state-of-the-art outlier detection tools such as OUTRIDER, OutSingle and FRASER in terms of computational speed and scalability. More importantly, saseR also dramatically boosts the performance for aberrant splicing (cf. FRASER) while maintaining a similar performance for aberrant expression detection (cf. OUTRIDER, OutSingle). Finally, we illustrate how utilizing these adapted offsets in conventional bulk RNA-seq tools allows to detect differential usage with superior computational time and similar detection performance (cf. DEXSeq), enabling the user to use a single tool for both DGE and DTU analyses.

## 2 Results

We first introduce how using adapted offsets can unlock conventional DGE frameworks for DTU analyses. We then elaborate on the extensions implemented in saseR to enable fast and scalable workflows for inferring aberrant expression or splicing. Next, we will use saseR to detect simulated aberrant splicing events, using adapted offsets, fast parameter estimation and count-based outlier tests. Then, we benchmark saseR for the detection of aberrant expression, and further showcase the use of adapted offsets in conventional bulk RNA-seq tools edgeR [22] and DESeq2 [23] to unlock these for differential transcript usage (DTU) analysis. Finally, we assess the performance of saseR to detect aberrant events in a case study with patients that have a validated disease mechanism.

### 2.1 Extending bulk RNA-seq tools for differential usage, aberrant expression and aberrant splicing

We start by introducing the modeling framework of conventional bulk RNA-seq tools and show how these can be extended for modeling differential usage and subsequently for detecting aberrant expression and splicing.

#### 2.1.1 Conventional bulk RNA-seq tools

Conventional bulk RNA-seq tools for differential gene expression analysis use a negative binomial framework to estimate the mean expression for each feature (e.g. gene expression read-counts, transcript expression read-counts) [22, 23] which can be formulated as:

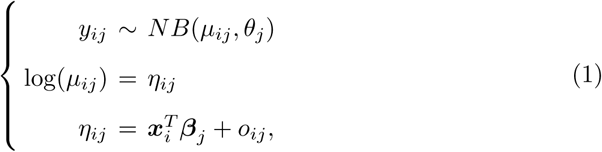

with *y_ij_* the observed count for feature *j* of sample *i*, *µ_ij_* the sample specific mean of feature *j*, *θ_j_* the negative binomial dispersion parameter for feature *j*, *η_ij_* the linear predictor of feature *j* for sample *i*, 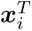 the covariate vector for sample *i*, ***β***_*j*_ a vector with the corresponding model parameters for feature *j*, and *o_ij_* an offset for feature *j* in sample *i* to normalise for differences in library size between samples. Note that conventional bulk RNA-seq tools by default consider the same offset for all features.

Changing the input features from genes to transcripts (or exons) enables bulk RNA-seq tools to assess differential transcript (or exon) expression. DEXSeq [24] further extends the model to accommodate for differential usage by modelling the feature (transcript or exon) counts together with the other counts for all features of the same gene in each sample. This approach implies the inclusion of a subject specific blocking covariate to address the correlation between the feature counts and the other counts in the same sample. The design matrix of DEXSeq thus grows with an additional dummy variable for each additional subject that is included in the study, which is leading to a huge computational burden when applying the tool to large data compendia.

#### 2.1.2 Changing offsets unlocks bulk RNA-seq tools for modelling proportions

Here, we will avoid the problem of including sample specific blocking covariates by specifying different offsets for each feature. We argue that similar results as DEXSeq can be obtained using a negative binomial model with an offset for the total count of all features that map to a gene. Indeed, the mean model parameters in Model (1) then also get an interpretation in terms of the log-ratio relative to the total count for a gene, which unlocks bulk RNA-seq tools for differential usage applications:

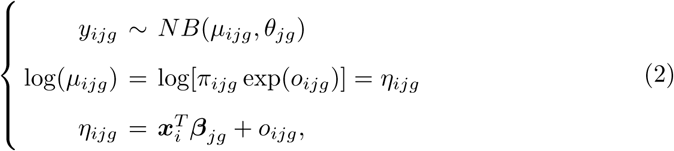

with *y_ijg_* the count for feature j of gene g in sample i, *π_ijg_* the usage for feature j of gene g in sample i, and 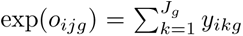 the total count over all features *J_g_* for gene g. We refer the reader to Supplementary materials for additional details on our rationale.

Note that an additional library size normalisation is not needed as the total gene count is subject specific. To avoid using the logarithm of 0 as offset, which occurs when the total gene count in a certain sample is 0, a pseudo-count of 1 is added to gene counts that are equal to 0, as well as to their corresponding feature count of 0. Upon parameter estimation, differentially used features (e.g. transcripts) are prioritised by testing on a single mean model parameter or a linear combination of model parameters that corresponds with the research hypothesis of interest. As conventional inference cannot be used for aberrant expression and splicing analyses, we will discuss count-based outlier tests in the next section.

Changing the offsets in the negative binomial framework to model proportions is generally applicable, and therefore allows us to extend bulk RNA-seq tools towards many different applications. Table 1 gives an overview for interesting applications that can be modeled with the bulk RNA-seq model by changing the input count table and offsets.

**Table 1.**
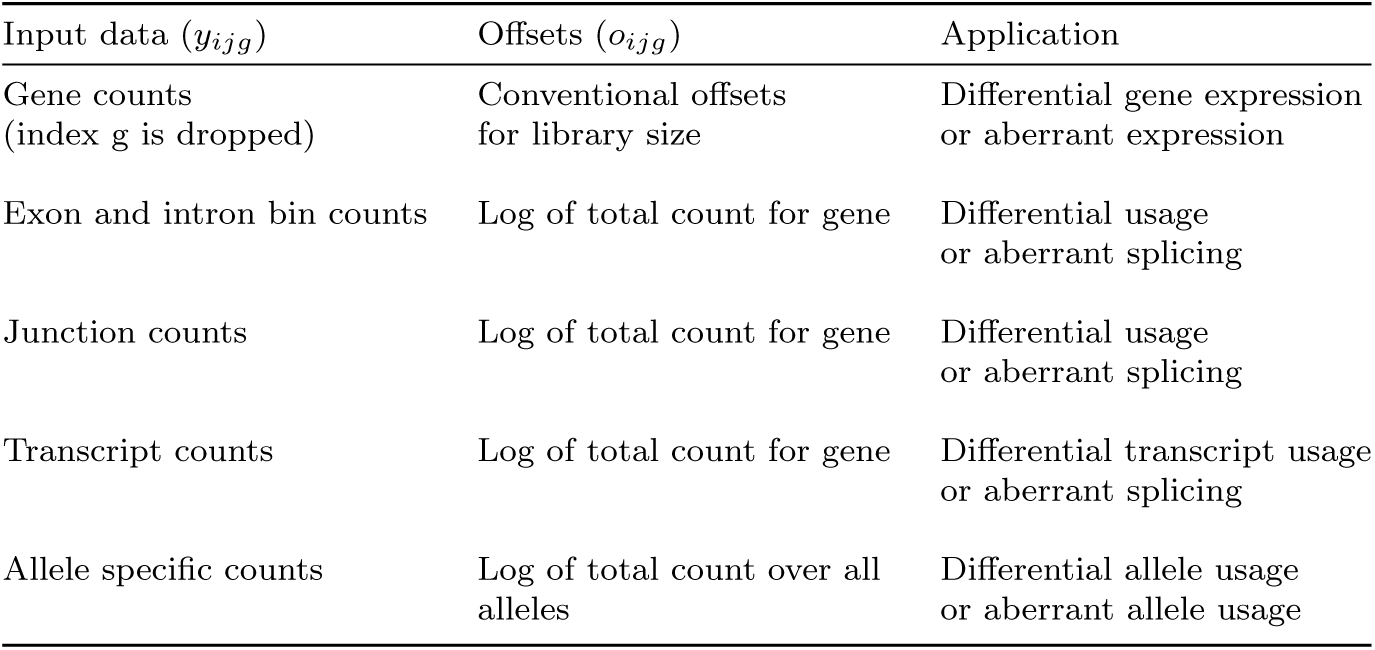
Bulk RNA-seq tools can be unlocked for different applications by carefully selecting the input count data and offsets for the negative binomial model framework.

| Input data ($y_{ijg}$) | Offsets ($o_{ijg}$) | Application |
| --- | --- | --- |
| Gene counts<br>(index $g$ is dropped) | Conventional offsets<br>for library size | Differential gene expression<br>or aberrant expression |
| Exon and intron bin counts | Log of total count for gene | Differential usage<br>or aberrant splicing |
| Junction counts | Log of total count for gene | Differential usage<br>or aberrant splicing |
| Transcript counts | Log of total count for gene | Differential transcript usage<br>or aberrant splicing |
| Allele specific counts | Log of total count over all<br>alleles | Differential allele usage<br>or aberrant allele usage |

For each application, one can use different counts and offsets in their workflow, and in particular workflows for intron and exon bin counts as well as junction counts are here developed for aberrant splicing. For the former we will use the logarithm of the total count over all bins that map to a gene as an offset. For the latter we will consider two workflows with different offsets, one with the logarithm of the sum over all junction counts that map to a gene, the other with an offset derived from ASpli [25] junction clusters, which do not require prior annotation. These junction clusters correspond to all junctions that have at least a splice site in common and are needed to infer novel/unknown splice sites.

#### 2.1.3 Count-based outlier test to detect aberrant expression and splicing

Conventional hypothesis testing for prioritisation of aberrant expression or splicing in the context of rare diseases is invalid because the disease-related features are different for each subject, and thus do not allow two-group comparisons due to the lack of replicates. Therefore, the problem changes to outlier detection. Indeed, aberrant features can be prioritised by plugging in the estimated mean model and dispersion parameters into the negative binomial distribution to obtain the corresponding quantile of the observed count for each feature in each sample. This quantile can be transformed to a kind of two-sided tail probability, which we refer to as a score in the remainder of the manuscript. Similarly to a p-value, small scores can be used for prioritizing features with outlying expression, and scores near one indicate a low evidence on outlying expression.

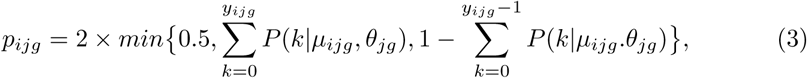

These can be estimated using conventional bulk RNA-seq tools such as edgeR [22]. Due to the discrete nature of the distribution, both tail probabilities can be greater than 0.5, requiring the restriction of the scores to be at most 1. Indeed, the lower and upper tail probability both include the count *y_ijg_*. These scores can be used to rank the genes according to their magnitude of aberrant expression or splicing. Note, that a distribution is needed to compute these quantiles, so we cannot resort to quasilikelihood based workflows like in satuRn [9] or the default quasi-likelihood workflow of edgeR.

Although the approach to use different offsets avoids many subject specific covariates in DTU or aberrant splicing analyses, the number of latent confounders that has to be controlled for in the latter applications is high because they rely on large data compendia, see e.g. state-of-the-art outlier detection algorithms OUTRIDER [1] and FRASER [20, 21]. Hence, negative binomial parameter estimation again becomes slow. We therefore implement a consistent parameter estimation algorithm in our saseR package that scales well to the large number of latent confounders typically included in aberrant expression and splicing analyses, which is possible because no inference has to be done on the parameter estimators in these applications (see Methods). Indeed, we only plug the mean model parameters in the NB distribution to calculate the corresponding quantiles. Further, saseR controls for latent confounders by using RUVr [27], and uses the Gavish-Donoho threshold [28] to select the appropriate number of latent confounders to be included in the outlier analysis (see Methods).

### 2.2 Detection of aberrant splicing

In this section we show how saseR unlocks the conventional negative binomial framework for fast and scalable aberrant splicing detection based on intron, exon and junction counts. Indeed, when using the logarithm of the total gene count as an offset, the mean model parameters again get an interpretation in terms of usages. To benchmark the performance of our workflows, aberrant splicing outliers are simulated in the lymphoblastoid cell lines from 39 healthy patients of the Geuvadis [29] dataset, using the RSEM simulator [30] (see Methods). The probability that splicing outliers are introduced in a gene of a particular patient was set at 0.001. Hence, this implies a different number of aberrant splicing outliers in a different set of genes for every subject. We evaluate different workflows for saseR: (1) saseR-bins that uses bin read counts (exon and intron) and the logarithm of the total gene counts as offset, (2) saseR-junctions relying on junction reads and the logarithm of the sum of the junction read counts per gene as offset, and (3) saseR-ASpli with junction reads and the logarithm of the sum of the junction reads that have at least one splice site in common, i.e. based on the ASpli junction cluster [25], as offset. These three saseR workflows are benchmarked against FRASER [20], which can use different autoencoders to control for confounders, i.e. a beta-binomial autoencoder (FRASER-Autoencoder), a PCA encoder and beta-binomial decoder (FRASER-BB-Decoder) and PCA (FRASER-PCA). All comparisons are based on the prioritisation of genes in which outliers were simulated. A gene-level score was obtained by using the minimal score of all features belonging to that gene.

Note that the results in the main paper are based on FRASER’s novel Intron Jaccard Index [21], which combines the former aberrant donor, acceptor and intron retention metrics [20]. We refer to this method as FRASER 2.0. The hyperparameter optimisation of FRASER was done with its PCA implementation to reduce computational time, which is its default setting. Results with FRASER’s donor and acceptor metrics are included in Supplementary Information.

Figure 2 shows the area under the precision-recall curve (AUC) for each sample, the overall precision-recall curve that considers the prioritisation in all samples together, and the computational time for saseR-bins, saseR-junctions, saseR-ASpli and FRASER 2.0. All saseR workflows outperform those of FRASER 2.0. saseR-bins and saseR-junctions have the best power, followed by saseR-ASpli. The precision-recall curve shows that the precision of FRASER 2.0 never reaches high levels, even at a low recall of simulated outliers. Further, saseR also outperforms LeafcutterMD [18] and SPOT [19] (Supplementary Fig. 1), which are tools that cannot control for known (e.g. batch effects) or latent confounder, typically present in clinical datasets.

**Fig. 2.**
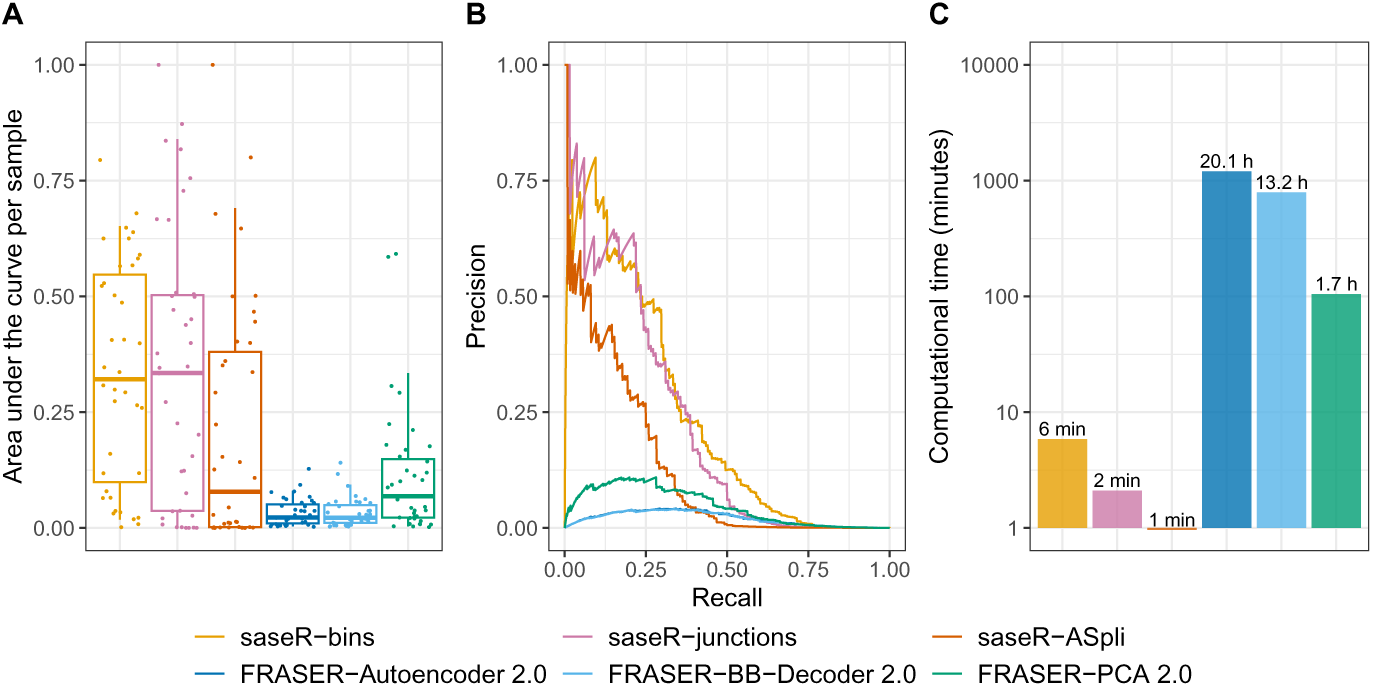
Benchmark of aberrant splicing detection. Comparison of performance to detect RSEM simulated splicing outliers in the Geuvadis dataset based on area under the precision-recall curve per sample (panel A), the precision-recall curve (panel B) and the computational time (panel C). saseR, with bin reads (saseR-bins), with junction reads and the logarithm of the total junction read counts per gene as offset (saseR-junctions), and with junction reads and the logarithm of the total junctions read count per ASpli junction cluster (saseR-ASpli) are benchmarked against FRASER 2.0 workflows, which use the Intronic Jaccard Index, i.e. one with an autoencoder (FRASER-Autoencoder 2.0), a beta-binomial decoder matrix (FRASER-BB-Decoder 2.0) and PCA (FRASER-PCA 2.0). Note, that PCA is always used for hyperparameter optimisation, which is FRASER 2.0’s default to reduce computational time. The whiskers of the boxplots in panel A correspond to the 5th and 95th quantile.

Remarkably, FRASER-PCA 2.0 performs better in our benchmark than its BB-Decoder and Autoencoder variants. This was not observed using the FRASER donor and acceptor metrics (Supplementary Fig. 2). But, the performance of these older methods never reaches that of FRASER-PCA, let alone that of saseR. To rule out convergence issues as the cause for the lack of performance of the FRASER 2.0 BB-decoder and autoencoder methods, we removed the junctions for which the decoder matrix did not converge. This, however, did not improve the results considerably (Supplementary Fig. 3).

The performances shown in Fig. 2 are solely based on the outliers that were included in its corresponding output, and filtered outliers are thus ignored. To ensure a fair comparison, we also assess the impact of the filtering strategies of saseR and FRASER. Alternatively, we assess the performance by enforcing all methods to use the same set of outliers. On the one hand, we consider the union of all outliers in the output of all methods. When an outlier is filtered for a specific workflow its score was set at 1. The results for this analysis remain very similar. The power of saseR-bins and FRASER remained similar. The power of saseR-junctions and saseR-ASpli reduced slightly because they filtered more outliers. However, they still largely outperform FRASER (Supplementary Fig. 4). On the other hand, we also considered the intersection of the outliers in all methods, which did not alter the results (Supplementary Fig. 5). The impact of the filtering strategy on the number of analysed outliers can be found in Supplementary Table 1.

Finally, we assess the performance when using FRASER’s injectOutliers function, starting with the option to simulate Jaccard outliers. For fair comparison, we used the junction read counts from the Kremer dataset on 119 fibroblast cell lines from patients diagnosed with mitochondrial diseases that were provided in the original FRASER paper [20]. saseR-junctions still largely outperforms FRASER-PCA 2.0, and FRASER with donor and acceptor metrics (Supplementary Fig. 6). saseR-bins, saseR-ASpli, LeafcutterMD and SPOT could not be benchmarked on the Kremer dataset, as only junction read-counts are publicly available. Note, that it is also possible to include junction counts and Jaccard offsets in saseR. This, however, does not lead to consistent results. It dramatically outperforms FRASER-PCA 2.0 in the Geuvadis benchmark and reaches similar performance as saseR-junctions (Supplementary Fig. 7), but has a slightly lower power in the Kremer benchmark compared to FRASER-PCA 2.0, which are both outperformed by saseR-junctions (Supplementary Fig. 8).

With FRASER’s injectOutliers function one can also simulate *ψ*_3_ or *ψ*_5_ outliers, which represent outlying usage of alternative donor or acceptor splice sites. Here, saseR-junctions’s performance remains similar as in other benchmarks, however, it is now slightly outperformed by FRASER-PCA workflows that specifically search for these *ψ*_3_ or *ψ*_5_ outliers (Supplementary Fig. 9 and 10). Interestingly, saseR can also infer outliers from *ψ*_3_ and *ψ*_5_ counts and their respective total counts, upon which it becomes on par with these FRASER-PCA workflows, again showing the advantage of saseR’s flexible framework.

Interestingly, saseR considering bins, junction or ASpli junctions is also much faster than FRASER. Indeed, by using the Gavish and Donoho threshold method [28] saseR by default does not require hyperparameter optimisation to select the number of latent factors (see Methods). saseR can also be run with similar hyperparameter optimisation as FRASER to select the number of latent confounders (Supplementary Fig. 11, Supplementary Information), while remaining much faster than FRASER.

Supplementary Fig. 12 also shows the benefits of saseR’s fast estimation procedure. Indeed, parameter estimation with edgeR [22] does not scale well with increasing number of latent factors or large design matrices. saseR for aberrant expression and splicing, however, by default considers a quadratic variance structure, which reduces each Newton-Raphson iteration to a matrix multiplication and thus scales well towards large design matrices (see Methods). Moreover, Supplementary Fig. 13 shows that our unbiased and fast parameter estimation procedure with 50 iterations is as good as the slower edgeR implementation. But, the additional speed gain of restricting the number of iterations to 1 or 3 cannot be justified as it seems to lead to a sub-optimal precision-recall.

### 2.3 Detection of aberrant expression

To benchmark aberrant expression detection, the GTEx [31] and Kremer [32] datasets are used. Only suprapubic skin cells were retained from the GTEx data, originating from 249 healthy deceased donors. Outliers were introduced in the gene expression count matrix at random with a probability of 0.001 (see Methods). Again, this implies a different number of outliers in a different set of genes for every subject. The performance of saseR is benchmarked against OutSingle [17] and OUTRIDER [1]. Similar to the default releases of OutSingle and OUTRIDER, saseR is run without controlling for known confounders. We include two OUTRIDER workflows, (1) OUTRIDER-Autoencoder using a negative binomial autoencoder to estimate and control for latent factors, and (2) OUTRIDER-PCA performing a principal component analysis on log-transformed counts, which is computationally more efficient but does not account for the count properties of the data.

Figure 3 shows the AUC for each sample, the overall precision-recall curve that considers the prioritisation in all samples together and the computational time on the GTEx data with simulated aberrant expression outliers. saseR, OutSingle and OUTRIDER-Autoencoder have a similar performance for detecting simulated aberrant expression outliers, and slightly outcompete OUTRIDER-PCA. saseR and OutSingle are much faster than OUTRIDER-Autoencoder and OUTRIDER-PCA, because the former by default does not require hyperparameter optimisation by using the Gavish and Donoho threshold [28]. Although OutSingle uses a log-normal approach, saseR is still twice as fast. saseR can also be run with a similar hyperparameter optimisation as OUTRIDER to select the number of latent factors (see Supplementary Fig. 14, Supplementary Information). This shows that, saseR with hyperparameter optimisation has similar performance compared to its fast default workflow, while remaining much faster than OUTRIDER-Autoencoder. Although saseR’s computational time with hyperparameter optimisation is slower than OUTRIDER-PCA, Supplementary Fig. 12 shows that the computational time to run a single analysis for a certain number of latent factors, genes and samples is faster for saseR. This indicates that the increased computational complexity is related to the outlier simulation scheme required for hyperparameter optimisation, i.e. NB versus Gaussian outliers, respectively.

**Fig. 3.**
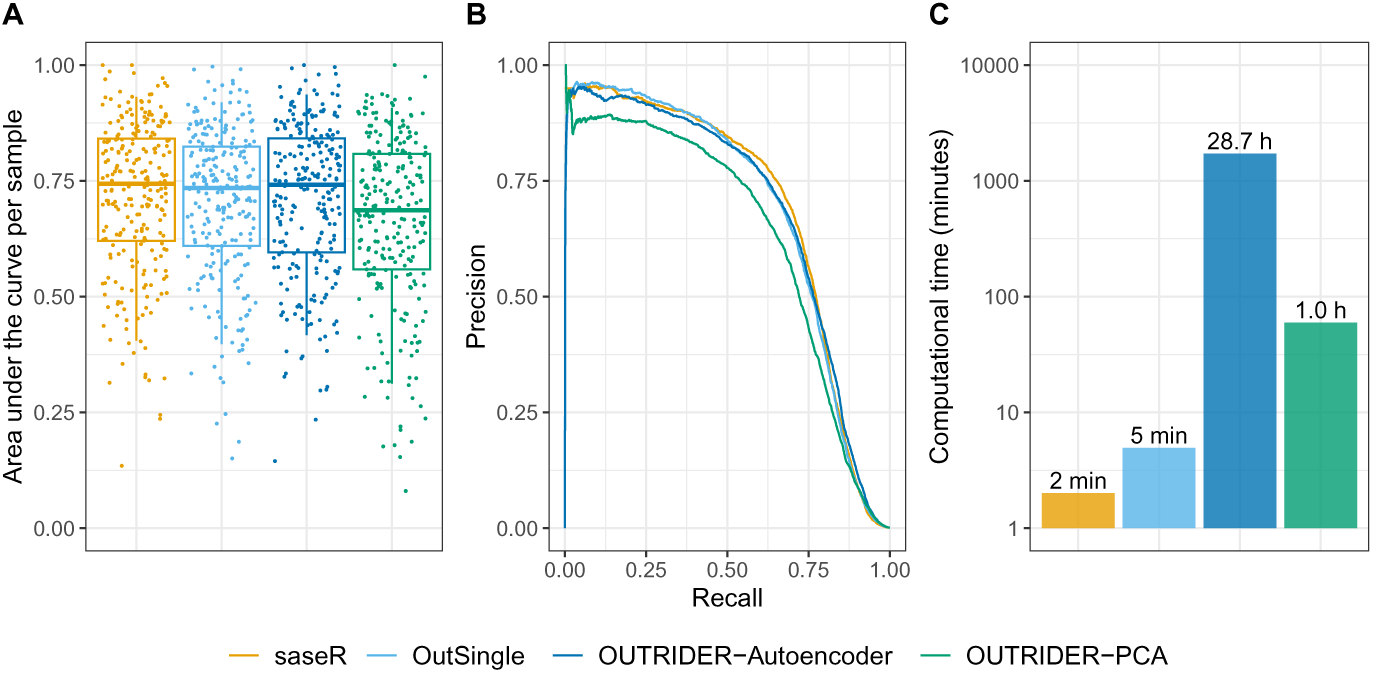
Benchmark of aberrant expression detection. Comparison of performance to detect simulated expression outliers in the GTEx dataset based on area under the precision-recall curve per sample (panel A), the precision-recall curve (panel B) and the computational time (panel C). Four methods are benchmarked: saseR, OutSingle, OUTRIDER-Autoencoder and OUTRIDER-PCA. Simulated outliers were introduced according to the gene-specific marginal distribution, only taking into account DESeq2 size factors for normalisation. The whiskers of the boxplots in panel A correspond to the 5th and 95th quantile.

Similar results are observed when analysing the Kremer dataset with simulated outliers (Supplementary Fig. 15).

Note, that saseR can also include known covariates to estimate the mean, although for the GTEx dataset this does not yield better performance to detect simulated outliers based on the conditional distribution (Supplementary Fig. 16). This functionality is not available for OutSingle.

### 2.4 Differential transcript usage analysis

Here, we show that the analysis of differential usage (DU), i.e. changes in relative abundance of transcripts/exons/introns within the same gene, can also be achieved with canonical bulk RNA-seq tools when using the logarithm of the total gene count as an offset. To assess the performance of these novel workflows, we add them to the benchmark of Gilis et al. [9], which assesses the performance to detect simulated DTU events in both bulk and single-cell RNA-seq (scRNA-seq) (see Methods). Panels A and B in Fig. 4 show the performance to pick up DTU using true positive rate (TPR) versus the false discovery rate (FDR) plots for both bulk- and scRNA-seq datasets, respectively. Panel C shows the computation time in function of the number of samples. DEXSeq, edgeR-diffspliceDGE, satuRn, and our novel edgeR and DESeq2 workflows with adapted offsets are included in our comparison. Note, that the computational time for DEXSeq is not included in panel C because it was several orders of magnitudes slower, but is shown in Supplementary Fig. 17. The computational time of our novel edgeR and DESeq2 workflows are in line with the other methods. The performances of edgeR and DESeq2 using adapted offsets to detect DTU on bulk RNA-seq data are comparable to DEXSeq and satuRn, and outperform edgeR-diffspliceDGE. On scRNA-seq data, however, satuRn still outperforms all other methods. DEXSeq performs slightly better than edgeR with adapted offsets, closely followed by edgeR-diffspliceDGE and DESeq2 with adapted offsets. Note, however, that this comparison only involved 20 vs 20 cells as DEXSeq does not scale to the data volumes in real scRNA-seq datasets (see Supplementary Fig. 17).

**Fig. 4.**
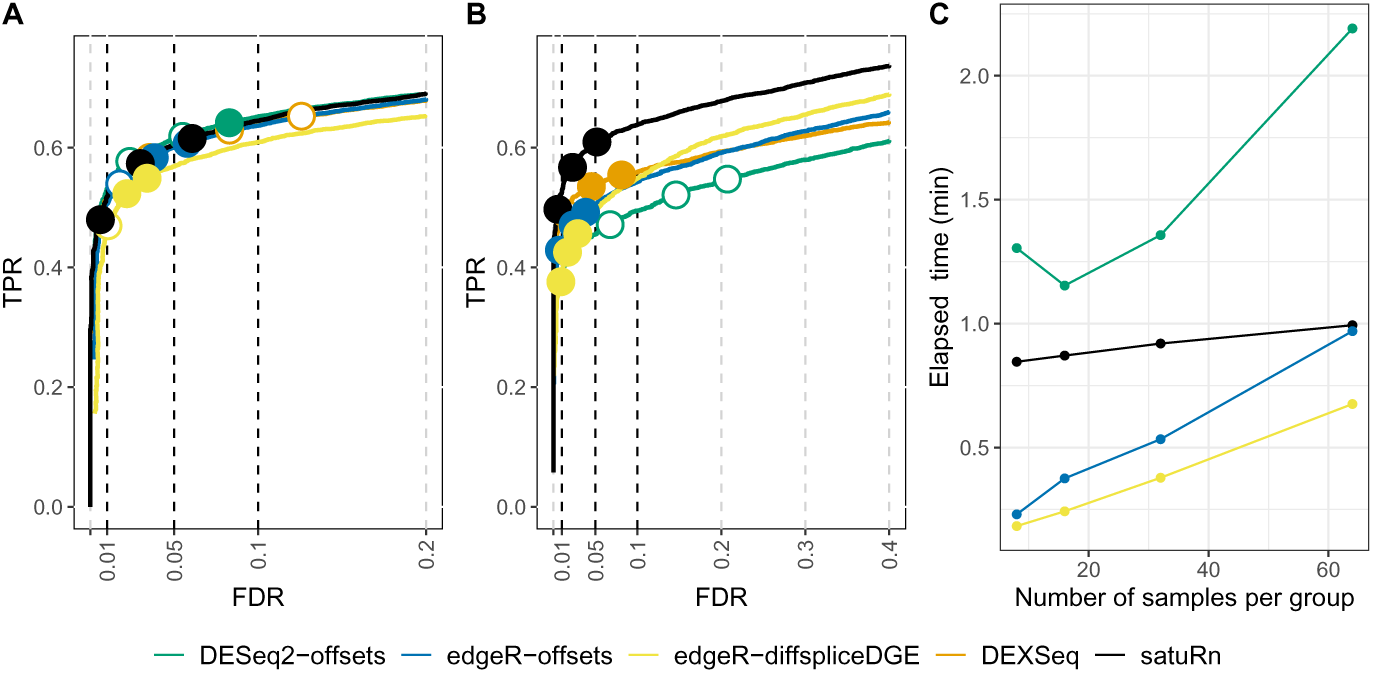
Benchmark of differential usage detection. Comparison of performance for differential transcript usage between satuRn, DEXSeq, edgeR-diffspliceDGE, and edgeR and DESeq2 with adapted offsets. The performance of the methods is compared on basis of true positive rate (TPR) versus false discovery rate (FDR) curves for a 5 vs 5 comparison on bulk RNA-seq data (panel A), for a 20 vs 20 comparison on scRNA-seq data (panel B) and the computational time relative to the number of samples (panel C). The three circles on each TPR-FDR curve represent the working points when the FDR level is set at nominal levels of 1%, 5% and 10%, respectively. The circles are filled if the empirical FDR is equal or below the imposed FDR threshold. Note, that the computational time of DEXSeq is not shown in panel C because it is several orders of magnitude larger than for the other tools, which was already illustrated in Supplementary Fig. 17.

### 2.5 Case study: Kremer dataset

saseR, OutSingle, OUTRIDER and FRASER are compared to detect aberrant expression and splicing events in rare disease cases from the Kremer dataset. This dataset includes 119 samples suspected to suffer from mitochrondrial diseases. However, the causal gene mutation is thought to differ between different patients, therefore requiring an outlier detection approach rather than a two-group comparison. We here only assess if the methods can identify the novel reported genes discussed by Kremer et al. [32], Brechtmann et al. [1] and Mertes et al. [20], as well as the list of disease related genes with aberrant splicing that are reported in the FRASER paper [20]. This means that 12 different disease-related genes are searched in 12 different patients using a single analysis that includes 107 other patients to improve the characterisation of the reference distribution for each gene. Although some gene variants (*ALDH18A1* and *MCOLN1* ) are known to be related to mono-allelic expression, these were picked up by Kremer et al. [32] and are also considered here. Also, *TIMMDC1* was reported to be the disease-related gene in two different patients, *MUC1365* and *MUC1344*, and is therefore included twice in our results. The rank of the score of the disease-related genes that were validated in a specific patient are shown in Table 2. Every line corresponds to a disease-related gene in a different patient, validated in literature. Again, only saseR-junctions is used for prioritising genes with aberrant splicing, because no BAM files are publicly available for the Kremer dataset.

**Table 2.**
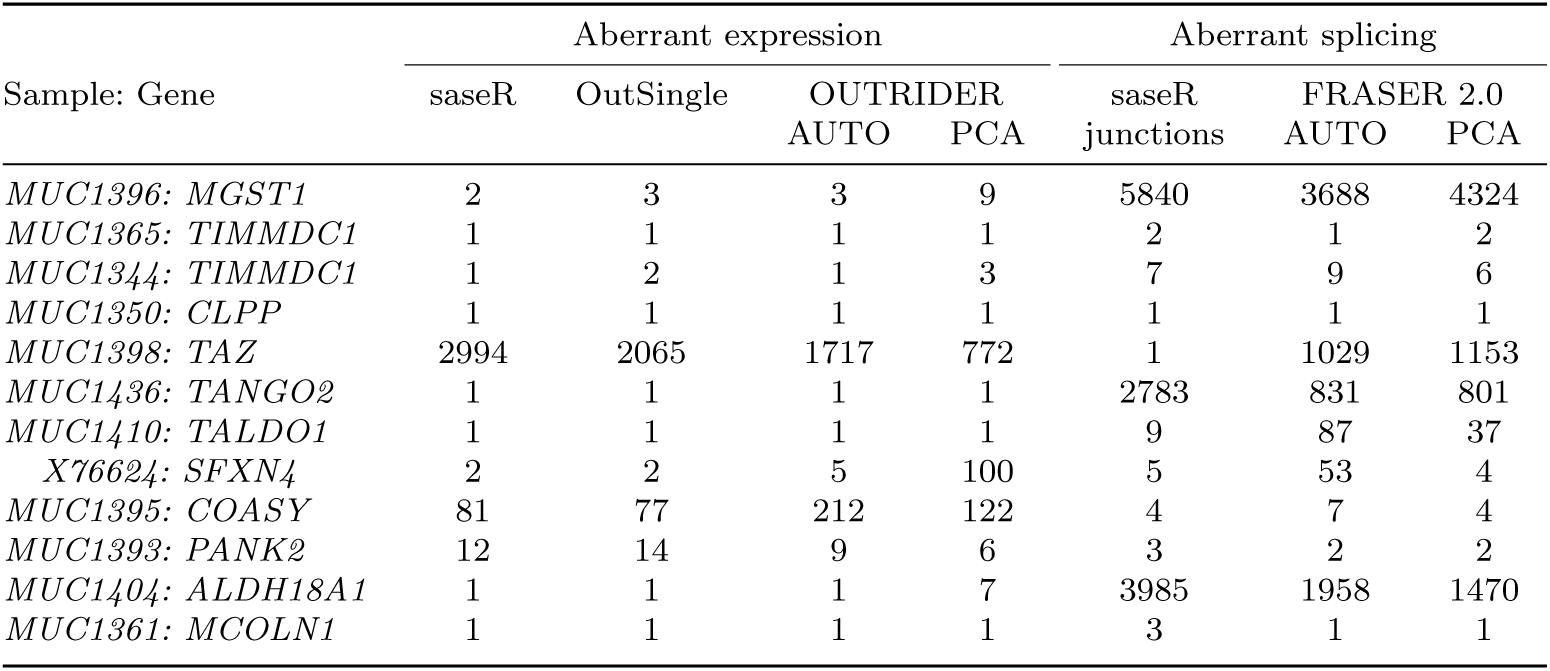
Detection of disease-related genes. Prioritisation based on the rank of the scores for disease-related gene within diagnosed patients using saseR, OutSingle, OUTRIDER AUTO (autoencoder), OUTRIDER PCA (principal component analysis), saseR-junctions, FRASER 2.0 AUTO and FRASER 2.0 PCA.

saseR aberrant expression, OutSingle and OUTRIDER show similar performance to prioritise the previously reported disease-related genes. Only *TAZ* and *COASY* are not easily detected by the four methods, while OUTRIDER-PCA additionally fails to prioritise *SFXN4*.

When assessing aberrant splicing with saseR-junctions and FRASER 2.0, it can be observed that the ranking of saseR-junctions is better in picking up the previously reported disease-related genes than FRASER 2.0 with the PCA and Autoencoder method. The most remarkable differences are *TAZ* and *TALDO1*, which are prioritised by saseR-junctions while they are missed by the FRASER 2.0 workflows. When using FRASER with donor and acceptor metrics (Supplementary Table 2) the prioritisation of most disease-related genes is worse. Also, a saseR workflow with junction counts and Jaccard offsets leads to suboptimal rankings compared to saseR-junctions and FRASER 2.0 (Supplementary Table 2).

Finally, an analysis with saseR using hyperparameter optimisation to determine the number of latent factors returns similar results as our default workflow with the Gavish and Donoho [28] threshold (Supplementary Table 3).

## 3 Discussion

In this contribution we developed saseR, a framework that unlocks bulk RNA-seq tools for fast and scalable differential splicing, aberrant splicing and expression analysis. Our key idea is to use specific RNA-seq counts in conjunction with well-chosen offsets to facilitate the proper interpretation of the mean model parameters for a specific application, i.e. gene counts and conventional offsets to correct for library size for the expression based analyses; transcript, exon, intron or junction counts with respectively the logarithm of the total count for all features per gene as an offset for usage and splicing based analysis; amongst others.

Upon parameter estimation, the conventional bulk RNA-seq inference framework can then be used to infer on differential expression or usage. When the aim is to infer aberrant expression or splicing, the estimated mean model and dispersion parameters are simply plugged into the negative binomial distribution to obtain the corresponding quantile to assess how extreme the observed count is for each feature in each sample.

Hence, our approach has the advantage of providing a single, unified framework to infer a wide range of applications. Moreover, in contrast to current state-of-the-art methods for aberrant splicing that only consider junction reads, it is also future-proof to novel sequencing-based technologies and applications, such as transcript counts with long-read sequencing and allele specific expression, amongst others, as long as the quantification can be recasted in specific feature counts in conjunction with proper offsets.

For aberrant splicing applications, saseR outperforms the current state-of-the-art method FRASER with donor, acceptor [20] and Jaccard metrics [21] both in terms of outlier detection as well as in computational complexity. saseR allows for different count inputs, such as bin read counts and junction read counts. This improves upon FRASER, which only uses junction read counts and discards much information on aberrant splicing that is also present in short-read RNA-seq exon and intron bin reads. With saseR we still provide a separate junction read count workflow, because bin read counts are less suited to pick up novel splice sites. We convincingly showed saseR’s superior performance in our simulation studies on aberrant transcript splicing outliers simulated with RSEM [30] as well as on junction outliers simulated with FRASER’s Jaccard outlier simulation scheme, and in a case study that focuses on prioritising disease-related genes in the Kremer dataset [32] that were reported in literature [1, 20, 32]. Note, however, that our benchmarks are purely based on transcriptomics data. Unlike Scheller et al. we did not have access to large-scale datasets with matched genomic data to provide orthogonal evidence in real applications.

For aberrant expression and differential usage on bulk RNA-seq data, the performance of the saseR workflows is at least on par with current state-of-the-art methods, such as OutSingle [17] and OUTRIDER [1] for aberrant expression, and DEXSeq [24] and satuRn [9] for differential usage analysis. However, saseR dramatically outperforms the existing methods in terms of computational time and/or flexibility to formulate the mean model.

The poor scalability of DEXSeq stems from modelling both the counts for a specific feature and the other counts for the same gene in one model, which requires the introduction of a dummy variable for each sample to address the within sample correlation. These sample specific intercepts, therefore, cause an explosion of the design matrix with increasing sample size. By normalising each feature with a well-chosen offset, e.g. the logarithm of the total feature count per gene in each sample, the mean model parameters also get the interpretation of a ratio without having to estimate a sample specific model parameter, which vastly improves the scalability. However, by using the total gene counts as a plugin estimator, their uncertainty is not propagated through the model. The same holds for the uncertainty on the transcript or exon bin counts due to multi-mapping reads, as these counts are also used as if they were observed.

For aberrant expression and aberrant splicing detection only an unbiased estimator for the mean model and dispersion parameters is required, which are subsequently plugged into the negative binomial distribution for outlier discovery. Therefore, saseR introduces a fast algorithm by assuming a quadratic variance structure when estimating the mean model parameters, which reduces each Newton-Raphson iteration to a matrix multiplication, which can be done simultaneously to update the parameters for all genes. This parameter estimator remains unbiased even when the variance structure is misspecified, which is known from generalized estimation equations theory [33]. We then use the mean model parameter estimates in edgeR’s *estimateDisp* function to estimate the negative binomial dispersion. This approach improves the scalability dramatically for applications on Mendelian diseases that rely on large cohorts and which typically require many latent factors to be included when estimating the mean model. We have shown that our fast algorithm for parameter estimation has equal performance to detect aberrant splicing events as our novel edgeR based workflows for usage applications, while improving the computational time by orders of magnitude for large studies. Our fast estimation approach, however, cannot be used in differential analyses because a misspecification of the variance structure renders the downstream inference invalid.

A further improvement upon OUTRIDER and FRASER is the use of Gavish and Donoho threshold [28] to determine the number of latent factors to include in the mean model, which avoids the computational intensive hyperparameter optimisation. Salkovic et al. [17] already introduced this idea in the context of aberrant expression detection. However, they considered RNA-seq read counts to be log-normally distributed, ignoring heteroscedasticity and the count nature of the data. Moreover, their method also lacks the flexibility to specify the mean model structure and cannot be used for other applications. We also implemented the option to select the number of latent factors in saseR with a similar hyperparameter optimisation as OUTRIDER and FRASER and showed that the performance for outlier detection remained very similar to the faster Gavish and Donoho threshold method.

## 4 Conclusion

We have developed a novel and very flexible framework, saseR, for fast and scalable analysis for differential usage, aberrant splicing and aberrant expression that dramatically outperforms state-of-the-art methods in terms of computational complexity. Interestingly, it also boosts the performance to detect aberrant splicing in rare diseases. Moreover, our approach has the advantage that it provides a unified workflow for many applications. Indeed, the user only has to change the input towards the proper count matrix and offsets for their specific application, which makes it generally applicable, user-friendly, and future-proof for current and novel sequencing-based technologies and applications. saseR is available on GitHub (https://github.com/statOmics/saseR/) and Bioconductor.

## 5 Methods

An introduction of the framework of saseR can be found in the Results section, explaining how it can infer aberrant expression, aberrant splicing and differential splicing using adapted offsets in the negative binomial framework. Here, we show how we control for unknown confounders, and develop a novel algorithm for parameter estimation that scales to large design matrices. We conclude with an overview of the datasets and our benchmarking protocols.

### 5.1 Correction for latent confounders

When performing differential or aberrant event detection in large data compendia one typically has to account for unknown confounders. To correct for these latent factors, several algorithms have been developed, e.g. [27, 34]. In saseR, we use the RUVr [27] approach. It first estimates the negative binomial model using known covariates to correct for treatment and batch effects. Then a singular value decomposition is done on the deviance residuals to estimate these latent confounders, which are subsequently incorporated in the negative binomial model as covariates. This approach computes the latent factors fast, while accounting for the heteroscedasticity in count data.

To estimate the optimal number of latent factors, two different approaches were used.

On the one hand, we implement the optimal hard threshold for singular values of Gavish and Donoho [28], which was adopted recently by Salkovic et al. [17] in the context of aberrant expression. The Gavish and Donoho threshold determines the optimal rank when denoising a matrix with Gaussian noise using a singular value decomposition. Salkovic et al. log-transformed the RNA-seq read counts, and performed both the matrix decomposition and outlier detection on the log-transformed scale to improve this Gaussian approximation. In contrary, RUVr performs this decomposition on the deviance residuals, which seems to work well in conjunction with the optimal number of latent factors by using the Gavish and Donoho threshold. These latent factors are then further included as covariates in the negative binomial regression model.

On the other hand, we also implement a similar approach to OUTRIDER [1] and FRASER [20, 21]: corrupted counts are simulated and replace the original counts with a frequency of 10*^−^*^2^. Then, a grid search is performed that varies the number of latent factors, and the prioritisation of the corrupted counts is then evaluated. The number of latent factors that obtains the highest area under the precision-recall curve is then used for the final analysis on the original data. Note, that a grid search implies that this approach will be much slower than the strategy from Gavish and Donoho [28] because the NB-model has to be fitted for each grid point. For details on the simulation of corrupted counts for aberrant expression and splicing, we refer the reader to Supplementary Information.

### 5.2 Fast parameter estimation for large design matrices

The mean model parameters ***β****_jg_* of the negative binomial model (2) are commonly estimated using a Newton-Raphson algorithm. It iteratively solves

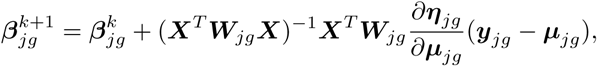

with ***W****_jg_* a *n* × *n* feature-specific diagonal weight matrix with elements

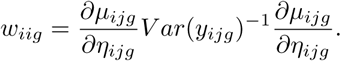

For the detection of aberrant events, and mainly for aberrant expression, the required number of latent factors to obtain optimal power can be large as large data compendia are typically used for this purpose. For large design matrices ***X***, the computation of the Newton-Raphson equation does not scale well and becomes slow. Therefore, we introduce a method for parameter estimation that is fast, scalable with large design matrices and builds on unbiased estimation equations, which are well known to provide consistent estimators (e.g. [35]). Particularly, we start from [36] who consider the class of unbiased estimation equations of the form

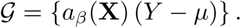

Lets define the functions *µ* and *a* as

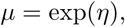

and

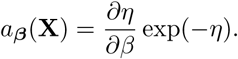

For a sample with *n* observations the estimating equations become

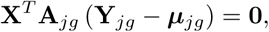

with **A***_jg_* a diagonal matrix with diagonal elements exp(−*η_ijg_*), which can be solved with the following Newton-Raphson algorithm:

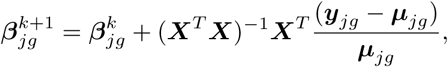

as the derivative of the estimating equations to ***β*** can be approximated by −**X***^T^* **X**.

Note, that these estimating equations and their corresponding Newton-Raphson algorithm also arise from Quasi likelihood upon replacing the NB variance structure, 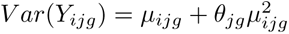 by a quadratic variance structure:

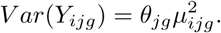

Indeed, the diagonal elements of weight matrix ***W****_jg_* in each Newton-Raphson iteration then reduce to a feature-specific constant, 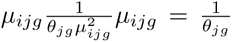, implying the Newton-Raphson update above.

Interestingly, the algorithm no longer involves feature-specific weight matrices, and shows that each iteration reduces to a fast matrix multiplication. Moreover, the part involving the design matrix in the update is the same for all features and their parameter estimates can thus be updated simultaneously in one large matrix operation.

Upon the estimation of the mean model parameters, the feature-specific dispersion *θ_j_* can be estimated using the functions of standard bulk RNA-seq tools, such as edgeR [22].

Note, that our fast parameter estimation can only be used for aberrant detection, which only requires an unbiased estimator of the mean and dispersion. Indeed, the estimators are then plugged into the NB distribution and no further inference is required on the parameter estimators themselves. Differential analysis, however, involves statistical hypothesis tests on (contrasts of) the mean model parameters, and the misspecification of the variance could lead to incorrect standard errors on (contrasts of) the mean model parameters and thus to incorrect inference for these applications. This could be addressed by replacing the variance covariance matrix of the mean model parameters by an appropriate sandwich estimator. However, that would again result in a computational burden and is not assessed, here. Finally, we assess bias and consistency of the implementation of our fast algorithm using simulations studies in Supplementary materials.

### 5.3 Data

Three different datasets are used in this work. First, similar to Brechtmann et al. [1], gene expression read counts were downloaded from GTEx portal (version V6P counted with RNA-SeQC v1.1.8) [31]. Only sequencing reads from suprapubic skin cells were retained and samples with a lower RNA integrity number than 5.7 were removed. Genes were kept when having at least 1 fragment per kilobase of transcript per million mapped reads for 5% of the samples. This filtering was done with DESeq2 [23]. Also, genes were only retained for analysis when having at least 1 or more read counts in 25% of the samples. This way, the analysed GTEx data contains 249 samples and 17065 genes. Second, gene, junction and intron-exon boundary reads were downloaded from Zenodo for the Kremer dataset [32, 37], which contains 119 samples suspected to suffer from Mendelian diseases. The gene expression read counts were filtered in the same way as the GTEx dataset, while the junction and intron-exon boundary reads were filtered using the standard filtering of FRASER [20]. This resulted in 14379 genes remaining for the aberrant expression analyses, and 79077 junctions remaining for the aberrant splicing analyses. For the case study, we considered searching the novel reported genes discussed by Kremer et al. [32], Brechtmann et al. [1] and Mertes et al [20], as well as the list of disease related genes with aberrant splicing that are reported in the FRASER paper [20]. Third, FASTQ-files of 39 samples from the Geuvadis dataset [29] were used (ERR188023-ERR188062, of which ERR188032 was removed due to errors with alignment), and were aligned to the protein coding genes of the of the GRCh38.p13 primary genome assembly [38] using STAR (version 2.7.10b) [39]. Bin and junctions read-counts were further filtered according to the default workflow of the corresponding tool (default edgeR [22] filtering for saseR).

### 5.4 Benchmarking simulations

We here discuss the different simulation frameworks used for benchmarking different tools to detect aberrant expression, aberrant splicing and differential usage.

#### 5.4.1 Aberrant expression

Different methods to detect aberrant expression were benchmarked by introducing simulated outliers in the GTEx and Kremer datasets. Outliers were introduced in the gene expression count matrix at random with a probability of 0.001. Hence, this implies a different number of outliers in a different set of genes for every subject. The value of the outlier was determined by using a quantile of the negative binomial distribution, specified by a gene-specific mean and dispersion parameter. These parameters are obtained by performing a negative binomial regression of the read counts with a linear predictor with only an intercept and an offset with sample specific size factors obtained by DESeq2 [23]. The quantile used in the benchmarks correspond to the quantile of Z=3 in the standard normal distribution. Both over- and underexpression outliers were simulated. This is conceptually similar to the outlier simulation scheme of OUTRIDER [1]. However, we use proper counts from a negative binomial instead of a log-normal approximation.

#### 5.4.2 Aberrant splicing

The probability that splicing outliers are introduced in a gene of a particular patient was set at 0.001. Again, this implies a different number of aberrant splicing outliers in a different set of genes for every subject. To simulate aberrantly spliced genes, the read counts corresponding to a transcript within that gene should be increased, while the read counts corresponding to another transcript should be decreased. As this cannot be done starting from a exon or junction count matrix, RSEM (version 1.3.0) [30] is used to simulate these outliers in a similar way as Soneson et al. [40] did for differential usage benchmarks. First, RSEM estimates transcripts per million expression values in each sample. Then, from randomly selected genes, the two most expressed transcripts were further used. Candidate transcripts were only considered if the total gene expression is greater than 100, the two transcripts both have an expression proportion larger than 10%, and the difference between both expression proportions is larger than 30%. The expected expression proportions of these two transcripts are then switched to simulate aberrant splicing outliers. Next, based on these expected proportions, a Dirichlet distribution is used to simulate new transcript counts per million. FASTQ-files were simulated based on these transcript counts per million, using the same library sizes as the original samples. These files were again aligned using STAR (version 2.7.10b) [39] to obtain exon and intron bin, and junction read count matrices that are further used as input for the algorithms.

Further, also FRASER’s [20, 21] outlier simulation framework was used for bench-marking aberrant splicing on the junction reads of the Kremer dataset [32]. This was done with the *injectOutliers* function, with which Jaccard outliers were simulated with a minimal proportion change of 20%.

#### 5.4.3 Differential splicing

The differential usage benchmarks were performed on a subset of the benchmark datasets from Gilis et al. [9]. There, the performance of the different DTU methods was evaluated on real bulk RNA-seq data. This was achieved by first subsampling a homogeneous set of ten samples from the large bulk RNA-seq dataset available from the GTEx consortium [31]. Subsequently, the ten samples were split arbitrarily in two groups, generating a comparison for which no differential usage between groups is expected. Finally, differential usage was artificially simulated in 15% of the genes. The number of differential used transcripts within a gene was determined with the simulation strategy of Van den Berge et al. [41]. For the selected transcripts, the counts of transcripts within the same gene are swapped in one group of samples, inducing DTU signal between the groups. The analysis was performed three times, each time with a different assignment of samples to groups and a different selection of genes with DTU signal. In a similar vein, the performance of the different DTU methods was evaluated on real single-cell RNA-seq data [9]. There, a biologically homogeneous subset of 40 cells was obtained from Chen et al. [42]. Cells were randomly split into two groups, and DTU signal was introduced by swapping transcript counts within a gene in one group of cells. We refer to the original publication for more details [9]. Note, that the Benjamini-Hochberg method [43] was used to control the FDR across all transcripts, which is the default method to address multiple testing in all tools considered in our benchmark.

## 6 Declarations

### Ethics approval and consent to participate

Not applicable.

### Consent for publication

Not applicable.

### Availability of data and materials

No data were generated for this study. The GTEx v6p dataset is available through dbGaP (accession number: phs000424.v6.p1) at https://gtexportal.org/home. Gene, junctions, and intron-exon boundary read counts from the Kremer dataset [32] were downloaded from Zenodo (https://zenodo.org/record/4271599 [37]). FASTQ files from the Geuvadis dataset [29] were downloaded from https://www.ncbi.nlm.nih.gov/sra. The data from Gilis et al. [9], used for the differential usage benchmarking are available at https://doi.org/10.5281/zenodo.6826603 [44]. The scripts used to simulate aberrant expression and splicing, and the code to reproduce the analyses, figures and tables are available on our companion GitHub repository for this paper: https://github.com/statOmics/saseRPaper/. saseR with vignettes for the different workflows is available on GitHub (https://github.com/statOmics/saseR/) and Bioconductor.

### Author’s contributions

A.S. and L.C. conceived and designed the study. A.S. implemented the methods. A.S. and J.G. analyzed the data. A.S. and L.C. wrote the paper. E.D.B., J.G., M.V.H. and D.R. contributed to discussions and revisions of the initial draft.

## Acknowledgments

The Genotype-Tissue Expression (GTEx) Project was supported by the Common Fund of the Office of the Director of the National Institutes of Health, and by NCI, NHGRI, NHLBI, NIDA, NIMH, and NINDS. The data used for the analyses described in this manuscript were obtained from the GTEx Portal on 10/21/22 under accession number phs000424.v6.p1.

## Supplementary information

Supplementary information is provided.

