## Supplementary Information for "Juggling offsets unlocks RNA-seq tools for fast and Scalable differential usage, Aberrant Splicing and Expression Retrieval"

The Supplementary Information consists of Supplementary Methods that introduces the simulation framework for corrupted counts, Supplementary Figures and Supplementary Tables.

### Supplementary Methods

#### Simulation of corrupted counts for hyperparameter optimisation

saseR provides the option to determine the number of latent factors using hyperparameter optimisation.

First, a strategy similar to OUTRIDER [1] is used when the aim is to detect aberrant expression. OUTRIDER replaces the original counts by corrupted counts with a probability of 0.01. The corrupted counts are simulated by using the $e^z$ quantile of the log2-normal distribution parameterised by mean $\mu_{ij}$ and standard deviation $\sigma_{\mu_j}$:

$$\mu_{ij} = \log_2\big(\frac{k_{ij}}{s_i} + 1\big),$$
$$k^c_{ij} = round(s_i 2^{\mu_{ij} \pm e^z \sigma_{\mu_j}}),$$

with $\mu_{ij}$ the log-transformed and library size normalised reads for gene j from sample i, $k_{ij}$ the observed read counts for gene j from sample i, $s_i$ the DESeq2 sizefactor [2] for sample i, $k^c_{ij}$ the simulated corrupted count for gene j from sample i, z randomly simulated from a normal distribution with mean log(3) and standard deviation log(1.6) and $\sigma_{\mu_j}$ the standard deviation of the log-transformed, normalised reads for gene j. Because z is simulated for every corrupted count, the quantile is different for each replaced count. Note that on the one hand, the mean of the log2-normal distribution, $\mu_{ij}$, is obtained by using only the original count that will be replaced, and that on the other hand, the standard deviation, $\sigma_{\mu_j}$, is obtained from the gene-specific marginal distribution of the log2-transformed normalised counts. To obtain an integer count as corrupted count, the quantile of this log2-normal distribution is rounded to the closest integer.

For saseR, we adopt a similar approach, but utilise the corresponding quantile from a negative binomial distribution instead. Instead of using the log-transformed normalised count as the mean for a log-normal distribution, we now use the original count as mean of the negative binomial distribution. Further, we estimate the gene-specific overdispersion $\theta_j$ using the gene-specific marginal distribution, instead of the variance $\sigma_{\mu_j}$ used in the OUTRIDER simulation. This overdispersion parameter is estimated by edgeR [3]. Using this negative binomial distribution, we obtain corrupted counts by using a simulated quantile $e^z$, with z again simulated from a normal distribution with mean log(3) and standard deviation log(1.6).

Next, a strategy similar to [4, 5] is adopted when the focus is on aberrant splicing detection. Note, that aberrant splicing detection is based on modelling proportions, e.g. the read counts mapping to a junction (bin) on the total read count for the corresponding gene. Counts are replaced by corrupted counts with a frequency of $10^{-2}$. The corrupted counts are derived from corrupted proportions that are simulated by introducing a proportion change to a random feature j in sample i, i.e.

$$\psi_{ij}^c = \psi_{ij} + \Delta\psi_{ij}^c$$

with $\psi_{ij}^c$ the corrupted proportion for feature j in sample i, $\psi_{ij}$ the observed proportion for feature j in sample i and $\Delta\psi_{ij}^c$ the simulated proportion change.

$\Delta\psi^c_{ij}$ is simulated from a uniform distribution between 0.2 and the maximal possible proportion change ($\Delta\psi^{max}_{ij}$):

$$\Delta\psi^c_{ij} = \pm U(0.2, \Delta\psi^{max}_{ij}).$$

The maximal proportion change $\Delta\psi^{max}_{ij}$ is the original proportion of the feature itself ($\psi_{ij}$) when the proportion is reduced, i.e. when $\Delta\psi^c_{ij}$ is negative, and is $1 - \psi_{ij}$ for positive $\Delta\psi^c_{ij}$. If $\Delta\psi^{max}_{ij} < 0.2$, the sign of $\Delta\psi^c_{ij}$ is changed and $\Delta\psi^{max}_{ij}$ is again computed.

The proportion of the other features $l \neq j$ belonging to the same feature group (e.g. gene), which is defined by the denominator of the proportion, are then changed by $\Delta\psi_{il} = -\Delta\psi^c_{ij}\frac{\psi_{il}}{\psi_{ij}}$.

From these new proportions, new counts are simulated for all features of the feature group, using the multinomial distribution and the observed total count for the feature group.

#### Rationale of using adjusted offsets to model differential usage

In Section 2.1.2. of the main manuscript we argued that the use of specific offsets can unlock bulk RNA-seq tools for novel applications, e.g. for differential usage. Recall the model:

$$\begin{cases} y_{ijg} & \sim NB(\mu_{ijg}, \theta_{jg}) \\ \log(\mu_{ijg}) & = \log[\pi_{ijg}\exp(o_{ijg})] = \eta_{ijg} \\ \eta_{ijg} & = \boldsymbol{x}_i^T\boldsymbol{\beta}_{jg} + o_{ijg}, \end{cases} \tag{1}$$

with $y_{ijg}$ the count for feature $j$ of gene $g$ in sample $i$, $\pi_{ijg}$ the usage for feature $j$ of gene $g$ in sample $i$, $\mu_{ijg}$ the mean expression read counts for feature $j$ of gene $g$ in sample $i$, $\theta_{jg}$ the overdispersion for feature $j$ of gene $g$, $\exp(o_{ijg}) = \sum_{k=1}^{J_g} y_{ikg} = y_{i.g}$

the total count over all features $j = 1 \dots J_g$ for gene g, $x_i^T$ the covariate vector for sample $i$ and $\beta_{jg}$ a vector of corresponding model parameters for feature $j$ of gene $g$.

Our approach can be motivated as follows: (1) Shot noise in bulk RNA-seq is known to be Poisson distributed [6]. If we repeatedly would sequence the same sample, exon bin counts, transcript counts, junction and gene counts can thus be assumed to be Poisson distributed, i.e. $\mathrm{P}(\mu_{ijg})$. Hence, if we jointly consider all feature counts $j = 1 \dots J_g$ of a gene $g$ in sample $i$ and if we assume them to be independent, their total gene counts can also be assumed to be Poisson distributed $\mathrm{P}(\mu_{ig})$ with mean $\mu_{ig} = \sum_j \mu_{ijg}$. (2) If we now parameterize $\mu_{ijg}$ as $\mu_{ijg} = \pi_{ijg}\mu_{ig}$, $\pi_{ijg}$ gets the interpretation of a usage. Interestingly, when considering the model for all features $j = 1 \dots J_g$ jointly, this is equivalent to modeling these joint counts using a mixed multinomial-Poisson model where the number of multinomial trials is random and follows a Poisson distribution with mean $\mu_{ig}$ see e.g. Terza and Wilson [7]. (3) The mean total feature count per gene $\mu_{ig}$, however, is unknown and has to be estimated. Here, we propose to use the observed total gene count $\hat{\mu}_{ig} = y_{i.g}$ as a plugin estimator. Note that with this plugin estimator, one could get the impression that the model would allow for "forbidden counts" as it does not exclude counts that are larger than the observed total count $y_{i.g}$. However, from a repeated sampling perspective it would not be unexpected to observe a count $y^*_{ijg}$ in a new experiment for feature $j$ that is larger than the average total gene count $\mu_{ig}$ especially if the feature has a high usage (Note, that the same reasoning holds for the total gene count $y_{i.g}$ that was observed in the current experiment and is used as a plugin estimator for $\mu_{ig}$). Indeed, the confusion on "forbidden counts" simply arises by considering the counts as if they are generated from a multinomial with a fixed number of total trials $y_{i.g}$. (4) Sequencing data from multiple samples are typically overdispersed w.r.t. the Poisson distribution and are therefore commonly modeled using a negative binomial distribution. Interestingly, the pendant for the negative binomial distribution also exists: [8] for instance note that the sum of multiple negative binomial variables $y_{ijg} \sim NB(\mu_{ijg}, \theta_g)$ is also negative binomial distributed $y_{i.g} \sim NB(\mu_{ig}, \theta_g)$ provided that all features from the same gene share the same overdispersion parameter $\theta_g$. (5) Similar to DEXSeq we can relax these assumptions by modeling the counts for each transcript $j$ using a negative binomial with a separate dispersion parameter $\theta_{jg}$. Indeed, DEXSeq models the transcript counts $y_{ijg}$ and the other counts $y_{i.g} - y_{ijg}$ for each transcript jointly using a negative binomial with a separate overdispersion parameter $\theta_{jg}$. Note, however, that using our plugin estimator for the mean total gene counts $\mu_{ig}$ does not propagate its associated uncertainty in downstream statistical inference.

#### Bias in saseR's fast parameter estimation

Here, we asses the bias of saseR's fast parameter estimation when modeling gene expression counts in a realistic simulation study. Specifically, we start from filtered gene counts from the Kremer study and estimate the mean model parameters and dispersions with a conventional edgeR analysis. Next, we simulate data for each gene from a negative binomial distribution using these fitted model parameters and effective library sizes. We then estimate the mean model parameters both with edgeR as well as with our fast saseR estimation algorithm. Supplementary Figure 18 shows that both edgeR and saseR produce unbiased estimates and that their estimation errors are very similar.

#### Bias in usage estimation

In this section, we asses the bias in the usage estimation. Again, we simulate data by starting from the gene counts in the Kremer study. Specifically, we simulate data for a transcript $t$ and for the lumped counts of the other transcripts that we denote as $o$ by simulating negative binomial with $\mu_{itg} = \mu_{ij}\pi_{jg}$ and $\mu_{iog} = \mu_{ig}(1 - \pi_{jg})$, respectively. Note, that we used the estimated mean gene counts $\mu_{ig}$ and dispersions from the simulation in the Supplementary Section "Bias of saseR's fast parameter estimation", and, that the total observed feature count per gene than equals $y_{ig} = y_{itg} + y_{iog}$. In our simulations we also include results with alternative offsets: i.e the real average log-transformed gene count $\mu_{ig}$ which we refer to as the oracle offset, and model-based offsets obtained by modeling the simulated total gene counts using a conventional edgeR/saseR analysis to estimate $\mu_{ig}$.

In Supplementary Figure 19 we observe that an edgeR/saseR analysis with log transformed total counts as an offset only leads to a bias in the estimated proportions for features that have extremely high overdispersions. Particularly, features with low and high usages are affected. Interestingly, the bias disappears when using model-based offsets. Note, that variability on the estimator also seems to become very large when using the oracle offset for features with increasing usages. Interestingly, this does not seem to happen for edgeR and saseR approaches that rely on the observed total count or model-based offsets.

#### Calibration of two-sided tail probabilities

In this section, we employ the same simulation framework described in Supplementary Section: Bias in Usage Estimation. Specifically, for each simulated count, we compute the corresponding two-sided tail probability (1) under the model used to generate the data, i.e. with the means, usages and dispersions that were used to simulate the data; as well as with a negative Bionomial were we plugin the offsets, usages and dispersions estimated with different offset strategies: (2) the logarithm of the observed total feature counts, (3) the logarithm of the modeled total feature counts, and (4) the oracle offset. The results are presented in Supplementary Figure 20. The plot demonstrates that the two-sided tail probabilities are approximately uniform across a wide range of dispersions and usage levels. Deviations from uniformity are observed at low usage levels, and these occur across all methods, including those applied under the data-generating model. For large dispersions and/or high usage values ($\pi_{jg}$ greater than 0.8), however, the two-sided tail probabilities obtained using the total count offset show marked distortions, which suggests a reduced power to detect usage outliers for such features.

Importantly, in our practical applications, usage values are typically low; that is, bin and junction counts are generally small relative to their corresponding total feature counts. For example, in our analyses, only 0.24% of observed proportions exceeded 0.8 in the Geuvadis dataset, and only 0.31% in the Kremer dataset.

Our parametric simulations suggest that one might consider first modeling the observed total feature counts per sample with a conventional bulk RNA-seq analysis and then using their resulting log-transformed estimates as offsets, rather than relying directly on the observed total counts. However, across the more realistic simulations throughout our manuscript, we consistently observed lower performance with these model-based offsets, e.g. Supplementary Fig. 21. This is not surprising: as opposed to the parametric simulation in this section, for practical applications the true underlying structure of the total feature counts is unknown and can be misspecified. Fortunately, the observed total feature counts provide a more robust alternative towards such misspecification.

### Supplementary Figures

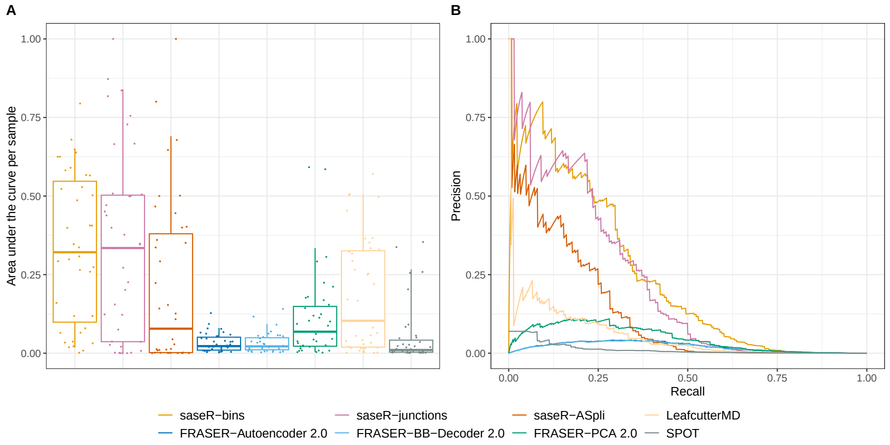


**Supplementary Fig. 1** Benchmark of aberrant splicing detection in the RSEM Geuvadis simulation. Comparison of performance based on area under the precision-recall curve per sample (panel A) and the precision-recall curve (panel B). saseR, with bin reads (saseR-bins), with junction reads and the logarithm of the total junction read counts per gene as offset (saseR-junctions), with junction reads and the logarithm of the total junctions read count per ASpli junction cluster (saseR-ASpli), are benchmarked to FRASER 2.0 [4, 5], LeafcutterMD [9] and SPOT [10] workflows. Note, that PCA is always used for hyperparameter optimisation for the FRASER workflows, which is FRASER 2.0's default to reduce computational time. The whiskers of the boxplots in panel A correspond to the 5th and 95th quantile.

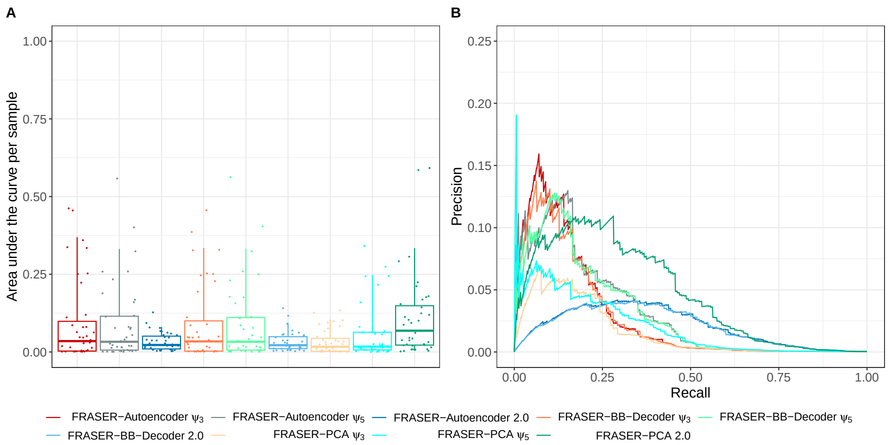


**Supplementary Fig. 2** Benchmark of aberrant splicing detection with FRASER. Comparison of performance to detect RSEM [11] simulated splicing outliers in the Geuvadis dataset [12] based on area under the precision-recall curve per sample (panel A) and the precision-recall curve (panel B). FRASER is benchmarked considering its three different metrics ($\psi_3$, $\psi_5$, Jaccard) and three different strategies to control for latent confounders (Autoencoder, BB-Decoder, PCA). Note, that PCA is always used for hyperparameter optimisation, which is FRASER's default to reduce computational time. The whiskers of the boxplots in panel A correspond to the 5th and 95th quantile.

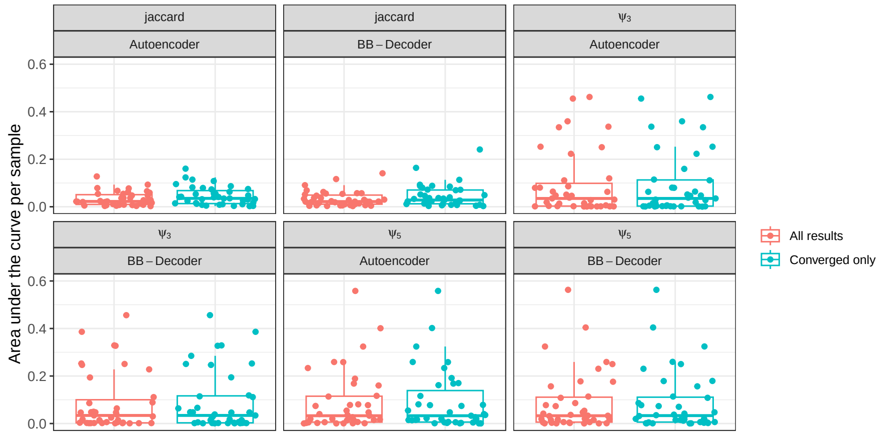


**Supplementary Fig. 3** Benchmark of aberrant splicing detection with FRASER while assessing the impact of convergence issues. The area under the precision-recall curve (AUC) per sample in the RSEM Geuvadis simulation is compared for FRASER considering its three different metrics ($\psi_3$, $\psi_5$, Jaccard) and two different methods to control for latent confounders (Autoencoder, BB-Decoder) that suffer from convergence issues. The AUCs are calculated based on the p-values as returned and based on p-values which are set at 1 if the features did not converge. Note, that PCA is always used for hyperparameter optimisation, which is FRASER's default to reduce computational time. The whiskers of the boxplots in panel A correspond to the 5th and 95th quantile.

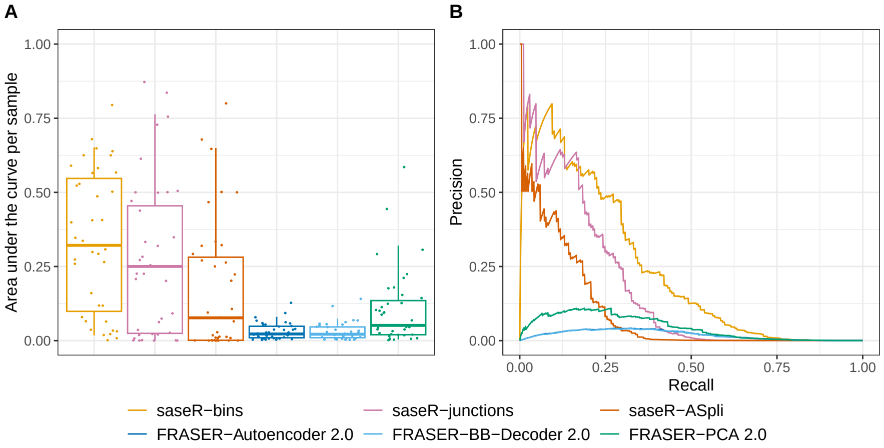


**Supplementary Fig. 4** Benchmark of aberrant splicing detection while accounting for filtered outliers. Comparison of performance in the RSEM Geuvadis simulation based on area under the precision-recall curve per sample (panel A) and the precision-recall curve (panel B). saseR, with bin reads (saseR-bins), with junction reads and the logarithm of the total junction read counts per gene as offset (saseR-junctions), and with junction reads and the logarithm of the total junctions read count per ASpli junction cluster (saseR-ASpli) are benchmarked to FRASER 2.0 workflows, which use the Intronic Jaccard Index and an autoencoder (FRASER-Autoencoder 2.0), a beta-binomial decoder matrix (FRASER-BB-Decoder 2.0) and PCA (FRASER-PCA 2.0). Outlier genes that were filtered by a method are added again and their p-value was set at 1 prior to the calculation of the performance metrics. Note, that PCA is always used for hyperparameter optimisation, which is FRASER 2.0's default to reduce computational time. The whiskers of the boxplots in panel A correspond to the 5th and 95th quantile.

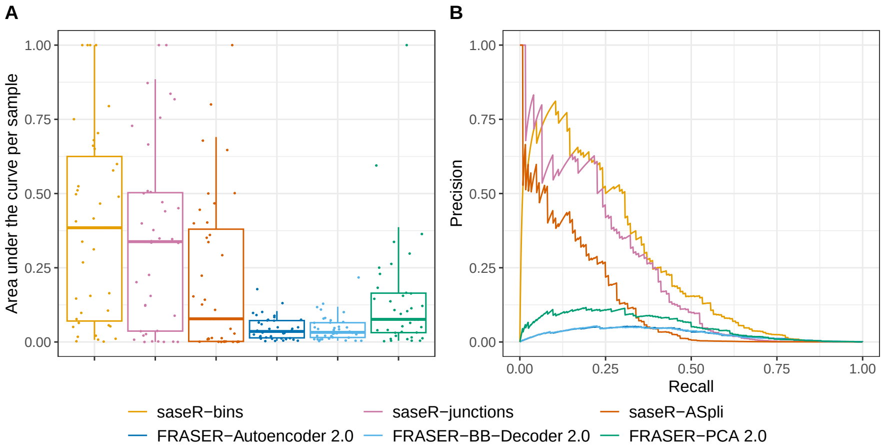


**Supplementary Fig. 5** Benchmark of aberrant splicing detection considering the intersection of returned genes. Comparison of performance in the RSEM Geuvadis simulation based on area under the precision-recall curve per sample (panel A) and the precision-recall curve (panel B). saseR, with bin reads (saseR-bins), with junction reads and the logarithm of the total junction read counts per gene as offset (saseR-junctions), and with junction reads and the logarithm of the total junctions read count per ASpli junction cluster (saseR-ASpli) are benchmarked to FRASER 2.0 workflows, which use the Intronic Jaccard Index with an autoencoder (FRASER-Autoencoder 2.0), a beta-binomial decoder matrix (FRASER-BB-Decoder 2.0) and PCA (FRASER-PCA 2.0). Only genes that were returned by all methods were retained. Note, that PCA is always used for hyperparameter optimisation, which is FRASER 2.0's default to reduce computational time. The whiskers of the boxplots in panel A correspond to the 5th and 95th quantile.

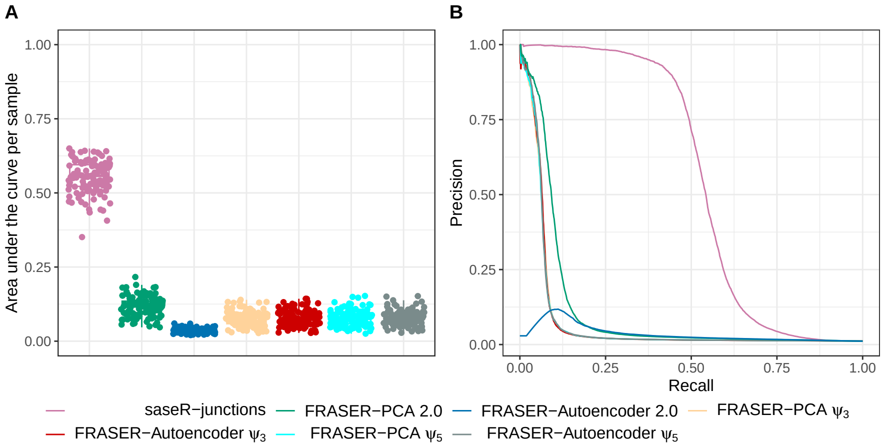


**Supplementary Fig. 6** Benchmark of aberrant splicing detection for simulated FRASER 2.0 jaccard outliers in the Kremer dataset. Comparison of performance based on area under the precision-recall curve per sample (panel A) and the precision-recall curve (panel B). saseR with junction reads and the logarithm of the total junction read counts per gene as offset (saseR-junctions) are benchmarked to FRASER workflows, which use the Intronic Jaccard Index (2.0), the donor ($\psi_3$) and acceptor ($\psi_5$) metrics, and consider an autoencoder (Autoencoder) or PCA to control for latent confounders. Outlier genes that were filtered by a method are added again and their p-value was set at 1 prior to the calculation of the performance metrics. Note, that PCA is always used for hyperparameter optimisation, which is FRASER's default to reduce computational time. The whiskers of the boxplots in panel A correspond to the 5th and 95th quantile.

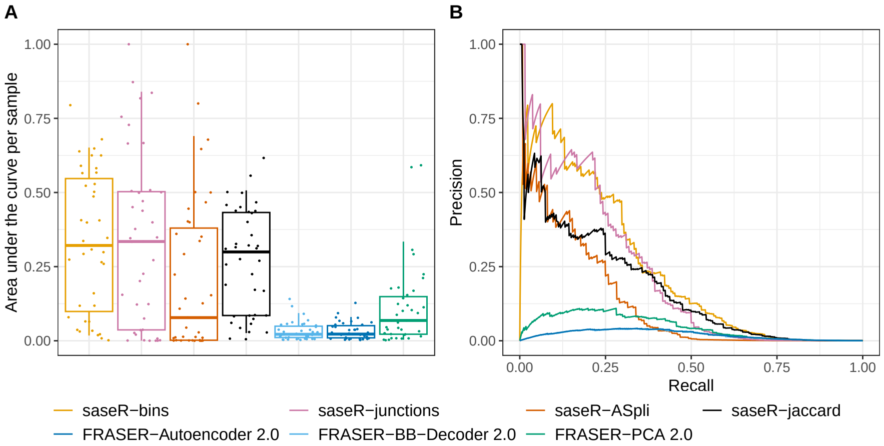


**Supplementary Fig. 7** Benchmark of aberrant splicing detection in the RSEM Geuvadis simulation. Comparison of performance based on area under the precision-recall curve per sample (panel A) and the precision-recall curve (panel B). saseR, with bin reads (saseR-bins), with junction reads and the logarithm of the total junction read counts per gene as offset (saseR-junctions), with junction reads and the logarithm of the total junctions read count per ASpli junction cluster (saseR-ASpli), and with jaccard counts and the sum of jaccard counts and other counts as offset (saseR-jaccard) are benchmarked to FRASER 2.0 workflows, which use the Intronic Jaccard Index with an autoencoder (FRASER-Autoencoder 2.0), a beta-binomial decoder matrix (FRASER-BB-Decoder 2.0) and PCA (FRASER-PCA 2.0). Note, that PCA is always used for hyperparameter optimisation, which is FRASER 2.0's default to reduce computational time. The whiskers of the boxplots in panel A correspond to the 5th and 95th quantile.

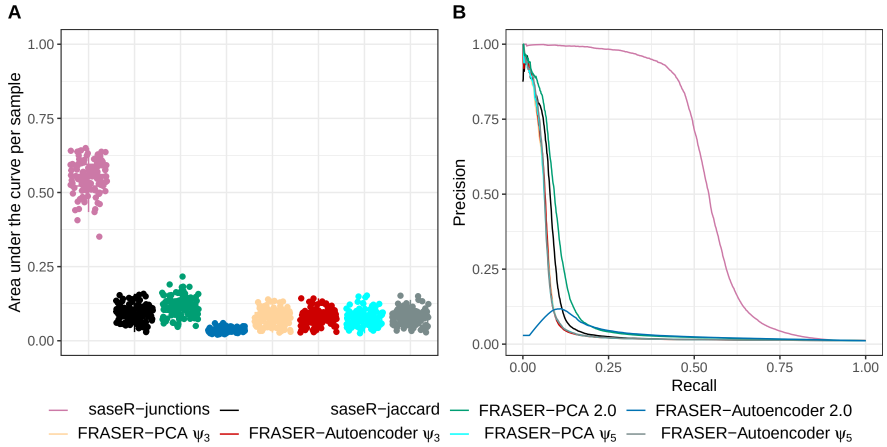


**Supplementary Fig. 8** Benchmark of aberrant splicing detection for simulated FRASER 2.0 jaccard outliers in the Kremer dataset. Comparison of performance based on area under the precision-recall curve per sample (panel A) and the precision-recall curve (panel B). saseR with junction reads and the logarithm of the total junction read counts per gene as offset (saseR-junctions), and with jaccard counts and the sum of jaccard counts and other counts as offset (saseR-jaccard) are benchmarked to FRASER workflows, which use the Intronic Jaccard Index (2.0), the donor ($\psi_3$) and acceptor ($\psi_5$) metrics, and consider an autoencoder (Autoencoder) or PCA to control for latent confounders. Outlier genes that were filtered by a method are added again and their p-value was set at 1 prior to the calculation of the performance metrics. Note, that PCA is always used for hyperparameter optimisation, which is FRASER's default to reduce computational time. The whiskers of the boxplots in panel A correspond to the 5th and 95th quantile.

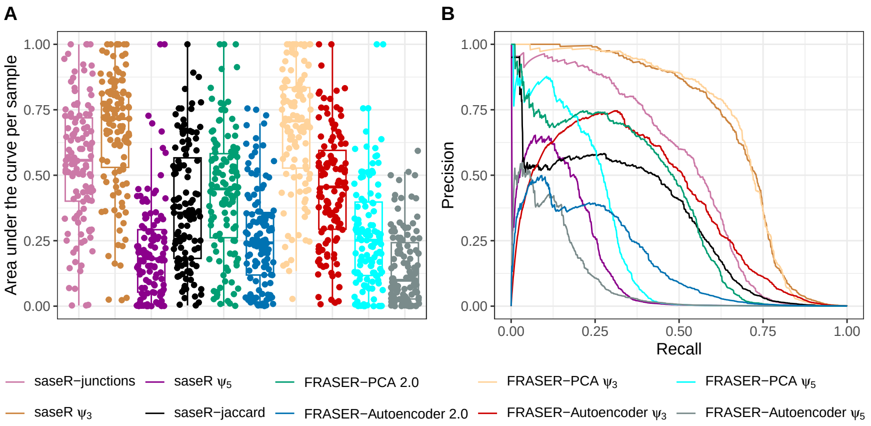


**Supplementary Fig. 9** Benchmark of aberrant splicing detection for simulated FRASER $\psi_3$ outliers in the Kremer dataset. Comparison of performance based on area under the precision-recall curve per sample (panel A) and the precision-recall curve (panel B). saseR with junction reads and the logarithm of the total junction read counts per gene as offset (saseR-junctions), with $\psi_3$ counts and the sum of $\psi_3$ counts and other counts as offset (saseR $\psi_3$), with $\psi_5$ counts and the sum of $\psi_5$ counts and other counts as offset (saseR $\psi_5$), and with jaccard counts and the sum of jaccard counts and other counts as offset (saseR-jaccard) are benchmarked to FRASER workflows, which use the Intronic Jaccard Index (2.0), the donor ($\psi_3$) and acceptor ($\psi_5$) metrics, and consider an autoencoder (Autoencoder) or PCA to control for latent confounders. Outlier genes that were filtered by a method are added again and their p-value was set at 1 prior to the calculation of the performance metrics. Note, that PCA is always used for hyperparameter optimisation, which is FRASER's default to reduce computational time. The whiskers of the boxplots in panel A correspond to the 5th and 95th quantile.

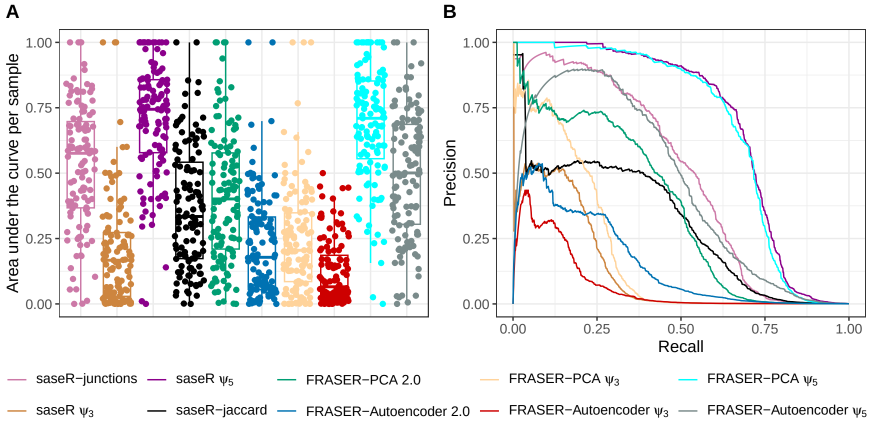


**Supplementary Fig. 10** Benchmark of aberrant splicing detection for simulated FRASER $\psi_5$ outliers in the Kremer dataset. Comparison of performance based on area under the precision-recall curve per sample (panel A) and the precision-recall curve (panel B). saseR with junction reads and the logarithm of the total junction read counts per gene as offset (saseR-junctions), with $\psi_3$ counts and the sum of $\psi_3$ counts and other counts as offset (saseR $\psi_3$), with $\psi_5$ counts and the sum of $\psi_5$ counts and other counts as offset (saseR $\psi_5$), and with jaccard counts and the sum of jaccard counts and other counts as offset (saseR-jaccard) are benchmarked to FRASER workflows, which use the Intronic Jaccard Index (2.0), the donor ($\psi_3$) and acceptor ($\psi_5$) metrics, and consider an autoencoder (Autoencoder) or PCA to control for latent confounders. Outlier genes that were filtered by a method are added again and their p-value was set at 1 prior to the calculation of the performance metrics. Note, that PCA is always used for hyperparameter optimisation, which is FRASER's default to reduce computational time. The whiskers of the boxplots in panel A correspond to the 5th and 95th quantile.

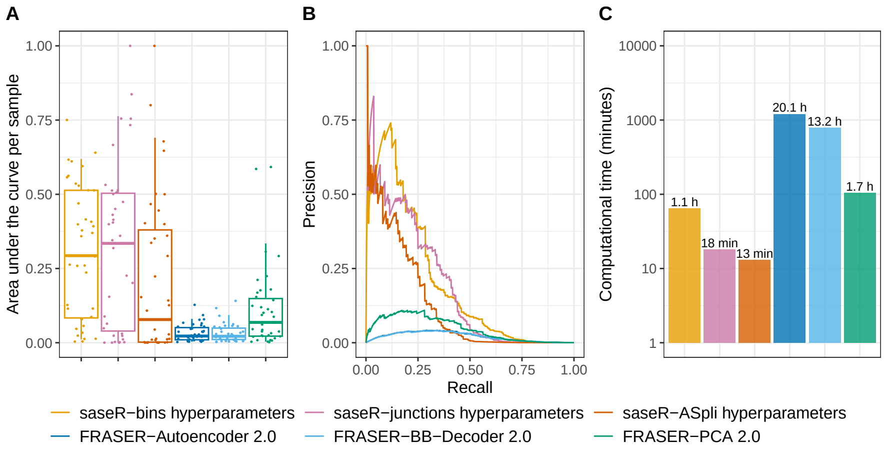


**Supplementary Fig. 11** Benchmark of aberrant splicing detection in the RSEM Geuvadis simulation. Comparison of performance based on area under the precision-recall curve per sample (panel A), the precision-recall curve (panel B) and the computational time (panel C). saseR using hyperparameter optimisation, with bin reads (saseR-bins), with junction reads and the logarithm of the total junction read counts per gene as offset (saseR-junctions), and with junction reads and the logarithm of the total junctions read count per ASpli junction cluster (saseR-ASpli) are benchmarked to FRASER 2.0 workflows, which use the Intronic Jaccard Index with an autoencoder (FRASER-Autoencoder 2.0), a beta-binomial decoder matrix (FRASER-BB-Decoder 2.0) and PCA (FRASER-PCA 2.0). Note, that PCA is always used for hyperparameter optimisation, which is FRASER 2.0's default to reduce computational time. The whiskers of the boxplots in panel A correspond to the 5th and 95th quantile.

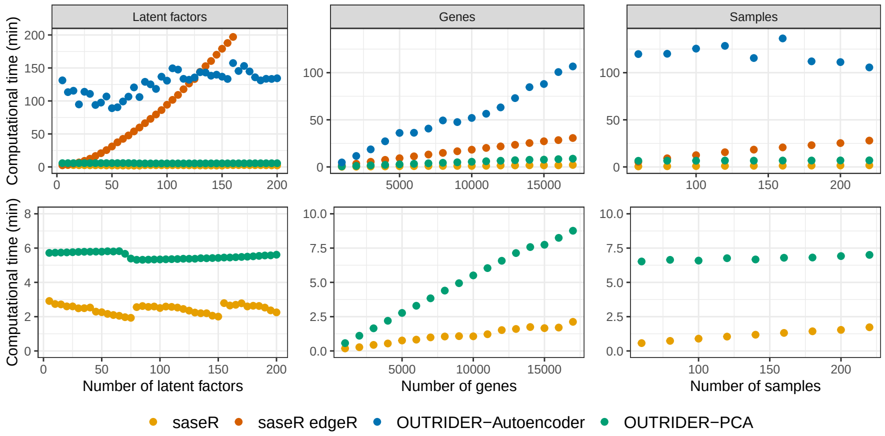


**Supplementary Fig. 12** Scalability benchmark of aberrant expression detection. Comparison of computational time to run an analysis of saseR, saseR edgeR, OUTRIDER-Autoencoder and OUTRIDER-PCA in the GTEx dataset at different number of latent factors, genes and samples. saseR edgeR uses edgeR [3] to estimate the mean model parameters. OutSingle is not included as it does not provide a workflow that is easily adapted. The left and middle plots use 249 samples, the left and right plots use 17065 genes, and the middle and right plots use 50 latent factors.

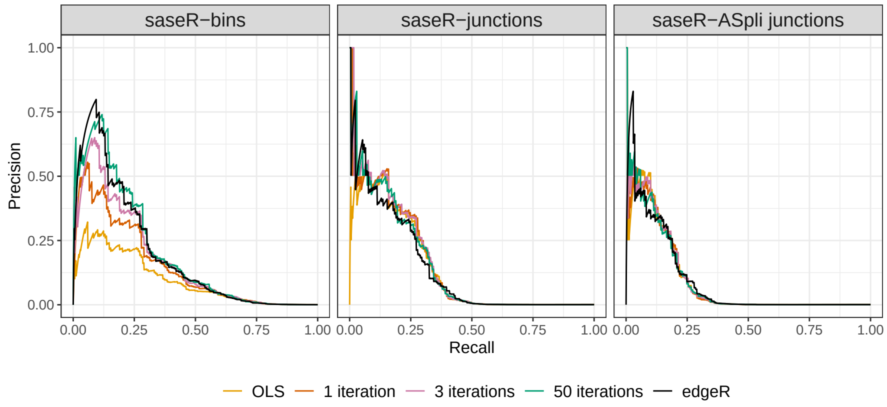


**Supplementary Fig. 13** Benchmark of aberrant splicing detection in the RSEM Geuvadis simulation. Comparison of performance based on the precision-recall curve. saseR, with bin reads (saseR-bins), with junction reads and the logarithm of the total junction read counts per gene as offset (saseR-junctions), and with junction reads and the logarithm of the total junctions read count per ASpli junction cluster (saseR-ASpli) are run with different parameter estimation methods. Our introduced parameter estimation with default iterations (50 iterations) is compared with ordinary least squares on log-transformed counts (OLS), restricting the number of iterations of our estimation procedure (1 iteration, 3 iterations) and edgeR. The same number of latent factors is used for each method.

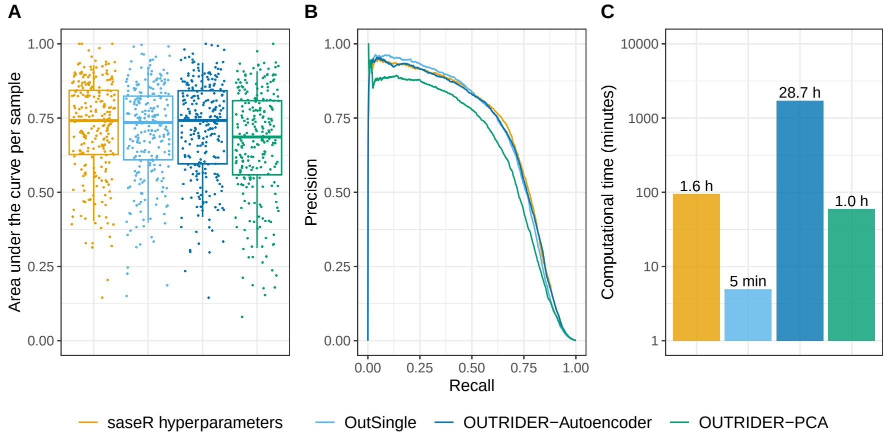


**Supplementary Fig. 14** Benchmark of aberrant expression detection. Comparison of performance to detect simulated expression outliers in the GTEx dataset based on area under the precision-recall curve per sample (panel A), the precision-recall curve (panel B) and the computational time (panel C). Four methods are benchmarked: saseR with hyperparameter optimisation, OutSingle [13], OUTRIDER-Autoencoder and OUTRIDER-PCA. Simulated outliers were injected according to the gene-specific marginal distribution, only taking into account DESeq2 size factors for normalisation. The whiskers of the boxplots in panel A correspond to the 5th and 95th quantile.

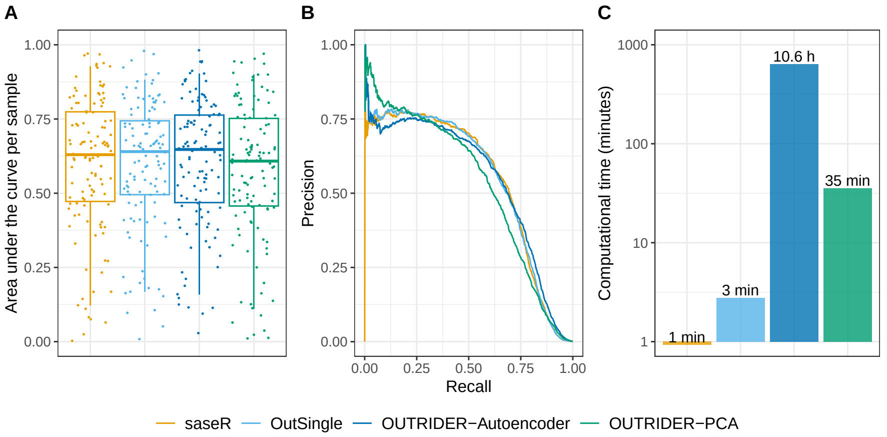


**Supplementary Fig. 15** Benchmark of aberrant expression detection. Comparison of performance to detect simulated expression outliers in the Kremer dataset [14] based on area under the precision-recall curve per sample (panel A), the precision-recall curve (panel B) and the computational time (panel C). Four methods are benchmarked: saseR, OutSingle, OUTRIDER-Autoencoder and OUTRIDER-PCA. Simulated outliers were injected according to the gene-specific marginal distribution, only taking into account DESeq2 size factors for normalisation. The whiskers of the boxplots in panel A correspond to the 5th and 95th quantile.

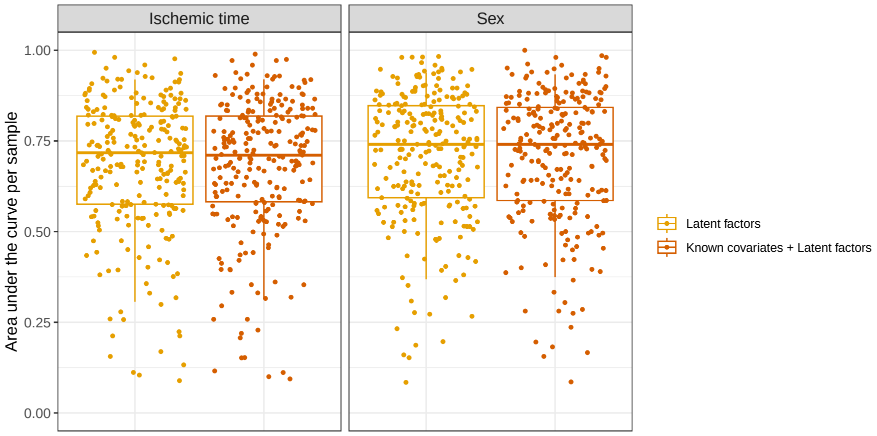


**Supplementary Fig. 16** Benchmark of aberrant expression detection with saseR when controlling for known covariates. The area under the precision-recall curve per sample is compared between saseR adding only latent factors and saseR that also includes known covariate Ischemic time (left) and Sex (right). Simulated outliers were injected in the GTEx dataset according to the gene-specific conditional distribution, considering the covariate Ischemic time (left) and Sex (right) together with DESeq2 size factors for normalisation. The whiskers of the boxplots correspond to the 5th and 95th quantile.

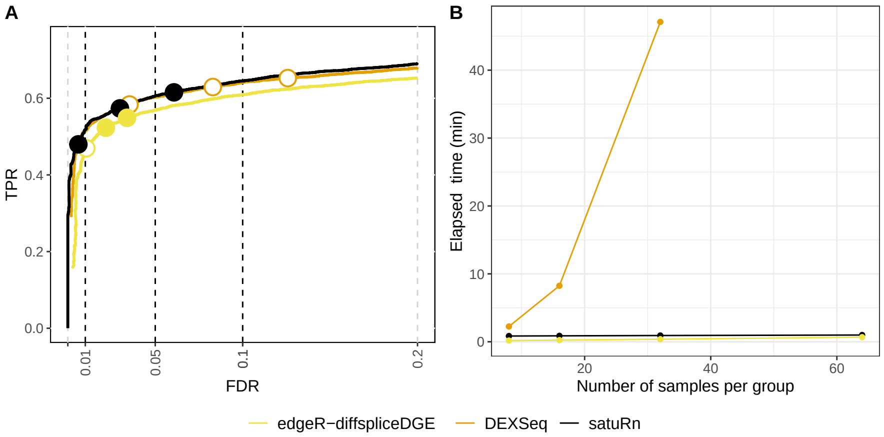


**Supplementary Fig. 17** Benchmark of differential usage detection. Comparison of performance for differential transcript usage between satuRn, DEXSeq and edgeR-diffspliceDGE. The performance of the methods is compared on basis of true positive rate (TPR) versus false discovery rate (FDR) curves for a 5 vs 5 comparison on bulk RNA-seq data (panel A) and the computational time relative to the number of samples (panel B). The three circles on each TPR-FDR curve represent the working points when the FDR level is set at nominal levels of 1%, 5% and 10%, respectively. The circles are filled if the empirical FDR is equal or below the imposed FDR threshold. The computational time of DEXSeq is several orders of magnitude larger than for the other tools.

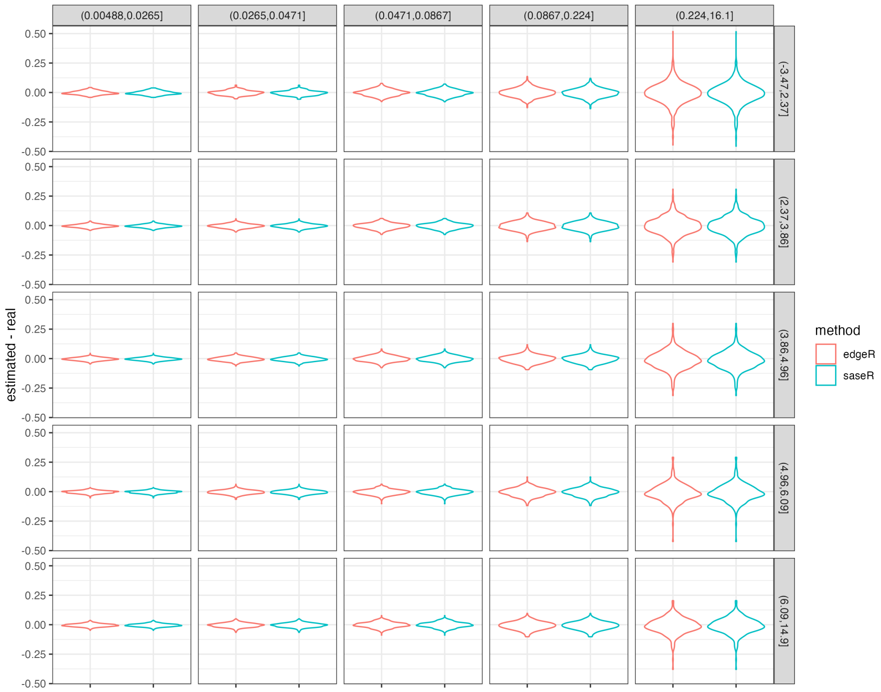


**Supplementary Fig. 18** Violin plots with the estimation error for the mean model parameters. Gene count data are simulated for 14379 genes and 119 samples with a negative binomial model using the dispersions, mean model parameters and effective library sizes estimated upon an edgeR on the Kremer data. In the y-axis the estimation error of the mean model parameters is plotted for edgeR and our fast saseR estimation. The panels are stratified according to the dispersion (horizontally) and the average log2 counts per million (vertically).

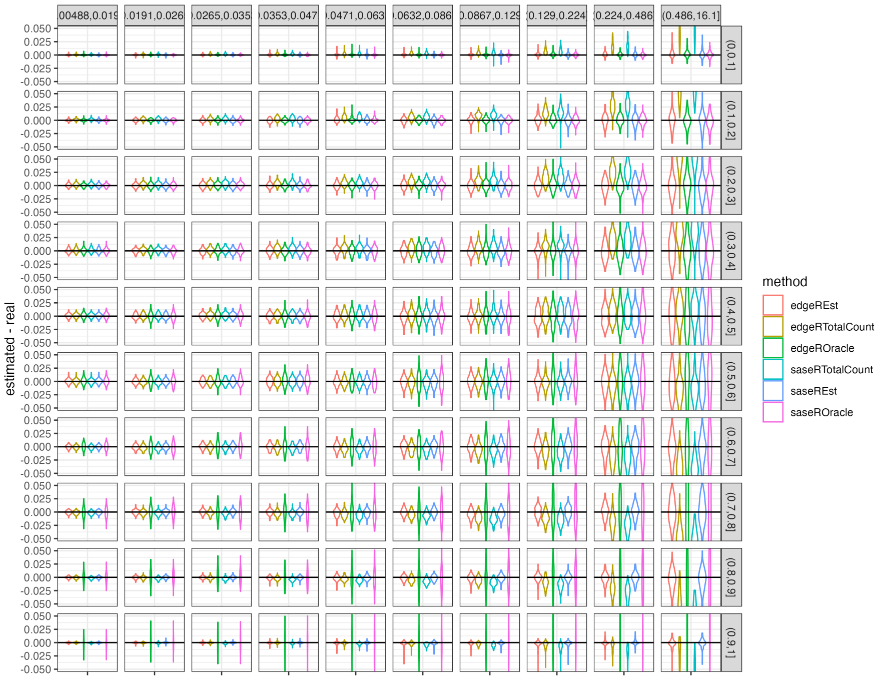


**Supplementary Fig. 19** Violin plots with the estimation error for usage. Usage count data are simulated with a negative binomial model using the estimated mean and dispersions from the Kremer data. The usages were simulated uniformly between 0 an 1 (for 14379 genes and 119 samples). In the y-axis the estimation error of the usage is plotted for edgeR and our fast saseR estimation using different offsets (Est: log-transformed average total feature count per sample obtained from an edgeR analysis on the observed total feature counts, Total Count: log-transformed observed total feature count and Oracle: log-transformed average gene count $\mu_{ig}$ that was used in the simulation). The panels are stratified according to the dispersion (horizontally) and the usage (vertically).

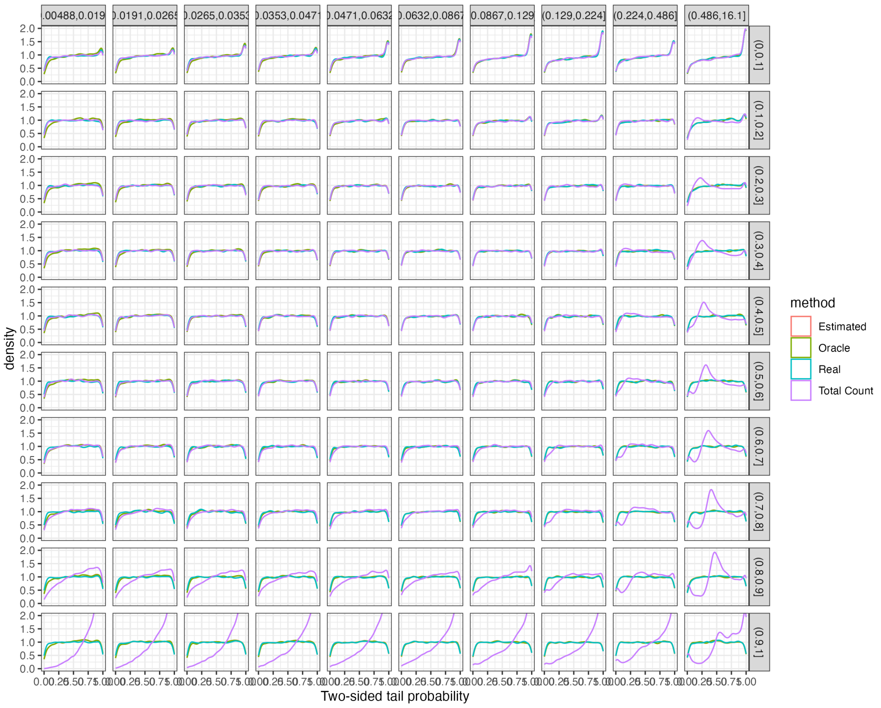


**Supplementary Fig. 20** Distribution of the two-sided tail probabilities. Usage count data are simulated with a negative binomial model using the estimated mean and dispersions from the Kremer data. The usages were simulated uniformly between 0 an 1 (for 14379 genes and 119 samples). The distribution of the two-sided tail probabilities are displayed as obtained with the parameters that were used for the simulation (Real) and upon estimation with negative binomial workflow using different offset strategies (Estimated: the log-transformed average total feature count obtained from an edgeR analysis on the observed total feature counts, Oracle: the log-tranformed average gene count $\mu_{ig}$ used in the simulation, and Total Count: the log-transformed observed total feature count). The panels are stratified according to the dispersion (horizontally) and the usage (vertically).

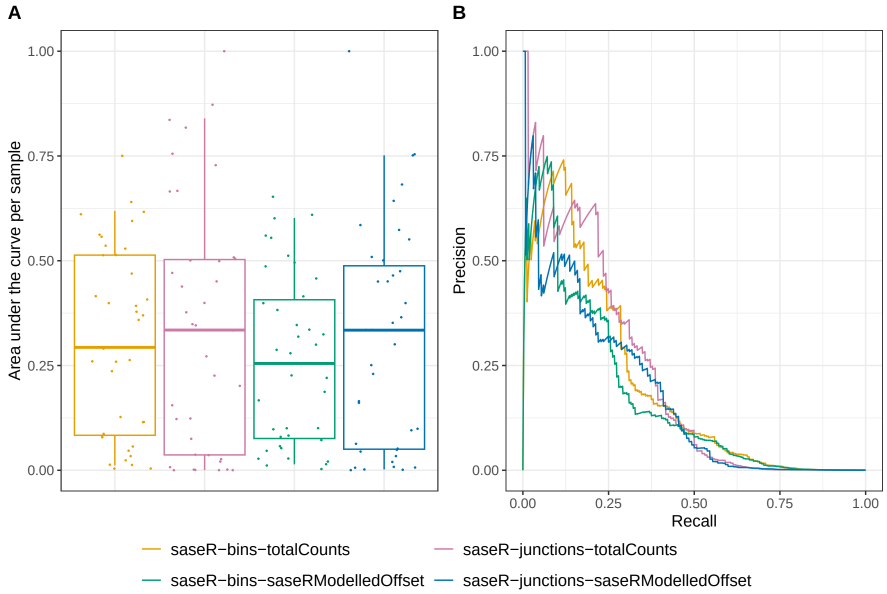


**Supplementary Fig. 21** Benchmark of aberrant splicing detection in the RSEM Geuvadis simulation. Comparison of performance based on area under the precision-recall curve per sample (panel A) and the precision-recall curve (panel B). saseR, with bin reads (saseR-bins) and with junction reads and the logarithm of the total junction read counts per gene as offset (saseR-junctions), are benchmarked with different offsets (totalCounts: the log-transformed observed total feature count, and modelledOffset: the log-tranformed average total feature count obtained by saseR estimation on the totalCounts level. The whiskers of the boxplots in panel A correspond to the 5th and 95th quantile.

### Supplementary Tables

**Supplementary Table 1** The number of outliers considered in the calculation of the performance criteria for the different filtering strategies used in the aberrant splicing analyses.

| | saseR | | | Fraser | |
|---|---|---|---|---|---|
| Analysis strategy | bins | junctions | ASpli | 2.0 | $\psi$ |
| Fig. 1: Considering only the returned genes of the method | 168 | 132 | 124 | 160 | 157 |
| Suppl. Fig. 4: Considering all the simulated outliers | 169 | 169 | 169 | 169 | 169 |
| Suppl. Fig. 5: Considering only the intersection of outliers | 124 | 124 | 124 | 124 | 124 |

**Supplementary Table 2** Detection of disease-related genes. Prioritisation based on the rank of the p-values for disease-related gene within diagnosed patients using saseR with junction reads and the logarithm of the total junction read counts per gene as offset (saseR-junctions), and with jaccard counts and the sum of jaccard counts and other counts as offset (saseR-jaccard), FRASER 2.0 AUTO (autoencoder), FRASER 2.0 PCA (principal component analysis), FRASER $\psi$ AUTO and FRASER $\psi$ PCA. Results of FRASER $\psi$ are merged from the donor ($\psi_3$) and acceptor metric ($\psi_5$) based on the best rank obtained for both metrics.

| | Aberrant splicing | | | | | |
|---|---|---|---|---|---|---|
| Sample: Gene | saseR | | FRASER 2.0 | | FRASER $\psi$ | |
| | junctions | jaccard | AUTO | PCA | AUTO | PCA |
| *MUC1396: MGST1* | 5840 | 3882 | 3688 | 4324 | 3223 | 3185 |
| *MUC1365: TIMMDC1* | 2 | 24 | 1 | 2 | 34 | 36 |
| *MUC1344: TIMMDC1* | 7 | 163 | 9 | 6 | 4 | 5 |
| *MUC1350: CLPP* | 1 | 1 | 1 | 1 | 1 | 1 |
| *MUC1398: TAZ* | 1 | 512 | 1029 | 1153 | 68 | 85 |
| *MUC1436: TANGO2* | 2783 | 349 | 831 | 801 | 128 | 157 |
| *MUC1410: TALDO1* | 9 | 6077 | 87 | 37 | 8598 | 1250 |
| *X76624: SFXN4* | 5 | 3 | 53 | 4 | 5741 | 8815 |
| *MUC1395: COASY* | 4 | 84 | 7 | 4 | 9496 | 9916 |
| *MUC1393: PANK2* | 3 | 3 | 2 | 2 | 1 | 1 |
| *MUC1404: ALDH18A1* | 3985 | 6820 | 1958 | 1470 | 5478 | 7042 |
| *MUC1361: MCOLN1* | 3 | 2 | 1 | 1 | 10738 | 11289 |

**Supplementary Table 3** Detection of disease-related genes. Prioritisation based on the rank of the p-values for disease-related gene within diagnosed patients using saseR, OutSingle, OUTRIDER AUTO (autoencoder), OUTRIDER PCA (principal component analysis), saseR-junctions, FRASER 2.0 AUTO and FRASER 2.0 PCA. saseR was run with hyperparameter optimisation.

| | Aberrant expression | | | | Aberrant splicing | | |
|---|---|---|---|---|---|---|---|
| Sample: Gene | saseR | OutSingle | OUTRIDER AUTO | OUTRIDER PCA | saseR junctions | FRASER 2.0 AUTO | FRASER 2.0 PCA |
| *MUC1396: MGST1* | 3 | 3 | 3 | 9 | 5576 | 3688 | 4324 |
| *MUC1365: TIMMDC1* | 1 | 1 | 1 | 1 | 1 | 1 | 2 |
| *MUC1344: TIMMDC1* | 1 | 2 | 1 | 3 | 7 | 9 | 6 |
| *MUC1350: CLPP* | 1 | 1 | 1 | 1 | 1 | 1 | 1 |
| *MUC1398: TAZ* | 1964 | 2065 | 1717 | 772 | 1 | 1029 | 1153 |
| *MUC1436: TANGO2* | 1 | 1 | 1 | 1 | 1041 | 831 | 801 |
| *MUC1410: TALDO1* | 1 | 1 | 1 | 1 | 5 | 87 | 37 |
| *X76624: SFXN4* | 7 | 2 | 5 | 100 | 7 | 53 | 4 |
| *MUC1395: COASY* | 351 | 77 | 212 | 122 | 4 | 7 | 4 |
| *MUC1393: PANK2* | 9 | 14 | 9 | 6 | 2 | 2 | 2 |
| *MUC1404: ALDH18A1* | 2 | 1 | 1 | 7 | 3376 | 1958 | 1470 |
| *MUC1361: MCOLN1* | 1 | 1 | 1 | 1 | 2 | 1 | 1 |
